# Solute carrier 5A5 regulates systemic glucose homeostasis by mediating glucose absorption in the *Drosophila* midgut

**DOI:** 10.1101/2021.06.10.447960

**Authors:** Yue Li, Weidong Wang, Hui-Ying Lim

## Abstract

The small intestine is the first organ that is exposed to and absorbs dietary glucose and thus represents the first of a continuum of events that modulates normal systemic glucose homeostasis. A better understanding of the regulation of intestinal glucose transporters is therefore pertinent to our efforts in curbing metabolic disorders. However, so far, the mechanisms known to regulate SGLT1, the primary intestinal glucose transporter, are mainly elucidated from in vitro studies. The *Drosophila* midgut, functional equivalence of the small intestine, could serve as an efficient in vivo model system for studying intestinal glucose transporter regulation; however, no glucose transporter has yet been identified in the midgut. Here, we report that the *Drosophila* Solute Carrier 5A5 (dSLC5A5) is homologous to SGLT1 and is highly expressed in the midgut. The knockdown of dSLC5A5 decreases systemic and circulating sugar levels and decreases glucose uptake into the enterocytes. In contrary, the overexpression of dSLC5A5 elevates systemic and circulating sugar levels and promotes glucose uptake into the enterocytes. We show that dSLC5A5 undergoes dynamin-dependent endocytosis in the enterocyte apical membrane, and that dSLC5A5 endocytosis is essential for the glucose uptake capability of dSLC5A5. Moreover, we provide evidence supporting that intracellular lysosomal degradation of endocytosed dSLC5A5 plays a significant role in the maintenance of dSLC5A5 level in the enterocyte apical membrane. We further show that short-term exposure to glucose upregulates SLC5A5 abundance in the enterocyte apical membrane. Finally, we show that the loss or gain of dSLC5A5 ameliorates or exacerbates the high sugar diet (HSD)-mediated glucose metabolic defects. Together, our studies uncovered the first *Drosophila* glucose transporter in the midgut and reveal new mechanisms that regulate glucose transporters in the enterocyte apical membranes.

## Introduction

The maintenance of normal systemic glucose homeostasis is vital for organismal fitness and viability. Upon its absorption from the intestinal lumen into the enterocyte, glucose travels transcellularly from the apical to the basal side of the cell and is secreted into the bloodstream to be transported to other organs. Glucose absorption in the small intestine is mediated by the Na^+^-glucose cotransporter SGLT1, which is enriched in the apical or brush border membrane of the mature enterocytes (Roder et al., 2014). SGLT1 is encoded by *SLC5A1*, the first of 11 gene members of the human solute carrier 5 (SLC5) family of plasma membrane Na^+^-dependent co-transporters that mediate the absorption of various solutes including sugars, inositol, vitamin, iodide and choline (Hummel et al., 2011; Wright et al., 1992).

The capacity for glucose absorption in the small intestine is tightly dependent on the availability of SGLT1, highlighting the importance of understanding the physiological regulation of the intestinal glucose transporter. To this far however, the mechanisms known to regulate SGLT1 are mainly elucidated from in vitro studies. Experiments performed in cultured cells and the *Xenopus* oocytes showed that SGLT1-containing vesicles undergo exocytosis from the trans-golgi network (TGN) to the plasma membrane, where SGLT1 is incorporated (Kroiss et al., 2006; Vernaleken et al., 2007; Veyhl et al., 2006). This intracellular trafficking pathway of SGLT1 is regulated by glucose, as shown by the observations that cellular injection of D-glucose or nonmetabolized glucose analogs accelerates the exocytosis of SGLT1 expressed in the *Xenopus* oocytes (Kroiss et al., 2006; Vernaleken et al., 2007; Veyhl et al., 2006). The incorporation of SGLT1 in the plasma membrane is balanced by the endocytosis of SGLT1 from the plasma membrane. SGLT1 has been documented to be endocytosed from the plasma membrane of cultured cells in a clathrin-and caveolin-independent manner (Ghezzi et al., 2017). Following endocytosis, SGLT1 could be targeted to the lysosomes for proteolysis or recycled back to the plasma membrane (Ghezzi et al., 2017). While these results have provided evidence supporting that endocytosis and the downstream lysosomal degradation and recycling are important intracellular trafficking pathways that regulate the availability of SGLT1 in the plasma membrane, they were obtained from in vitro work that utilized cultured cells which are not models for human enterocytes. Thus, it remained unclear whether the same mechanisms regulate glucose transporters in an in vivo setting. Furthermore, how the apical endocytosis of glucose transporters regulates glucose transporter-mediated glucose uptake in the small intestine is unclear. Finally, while glucose has been shown to regulate the exocytosis of SGLT1, it is not known whether endocytosis and the downstream recycling and lysosomal degradation of glucose transporters could be involved in the regulation of glucose transporter by glucose.

The *Drosophila* midgut is the functional equivalent of mammalian small intestine and could serve as an efficient in vivo model system to study the regulation of intestinal glucose transporter. In insects, glucose is the main fuel for energy metabolism, but is normally present at a low concentration in the hemolymph (insect blood) because of its reducing and osmotic properties (Becker et al., 1996). Glucose, once absorbed from the midgut [insect equivalence of mammalian small intestine (Lemaitre and Miguel-Aliaga, 2013)] into the hemolymph, is converted in the fat body into trehalose (Caccia et al., 2007). Trehalose, a disaccharide consisting of two molecules of glucose, is the dominant circulating sugar in *Drosophila* and represents a permanent source of glucose that can be continuously be released by enzymatic hydrolysis mediated by tissue-associated trehalases (Wegener et al., 2003). While sugars are critical substrates for insect metabolism, little is known about the transporters that mediate glucose absorption in the midgut. In *Drosophila*, up to a total of 78 *Drosophila* genes have been found to harbor a sugar transporter motif (Limmer et al., 2014). Therefore, it is likely that SGLT1-like glucose transporters exist in the midgut and play an essential role in mediating dietary glucose uptake. However, to this far, no dietary glucose transporter in the midgut has been reported. The identification of such transporters and the interrogation of their function and regulation in the *Drosophila* midgut will reveal new insights into the regulation of SGLT1 level and activity in the small intestine.

The *Drosophila* SLC5A (dSLC5A) family is homologous to the human SLC5 family of Na^+^-dependent solute transporters (Dus et al., 2013). Several members of the dSLC5A family have been characterized, including dSLC5A7 which is involved in salt tolerance (Stergiopoulos et al., 2009), dSLC5A11/cupcake which regulates feeding behavior (Park et al., 2016), and dSLC5A15 which serves as a choline transporter (Hamid et al., 2019). However, it is not known whether any member of the dSLC5A family functions as a glucose transporter in the midgut. Here, in this study, we report the identification of a member of the dSLC5A family, dSLC5A5, as the first dietary glucose transporter in the *Drosophila* midgut. We further reveal previously-unrecognized mechanisms that govern the abundance and activity of dSLC5A5/SGLT1 in the enterocyte apical membranes.

## Results

### Similarities between human SGLT1 and dSLC5A5

To identify the *Drosophila* glucose transporter in the midgut, we searched the dSLC5A family of genes using orthology data from the DIOPT dataset in FlyBase. Our analyses revealed a number of genes in the dSLC5A family that share close similarity (>40%) and identity (>20%) with human SGLT1 (hSGLT1). Among those members most highly conserved to hSGLT1, we performed qRT-PCR analysis of their endogenous expression in various tissues and identified dSLC5A5 that showed a high level of expression in the midgut amongst other tissues, including the skeletal muscle, fat body (functional equivalence of liver and adipose tissue), and brain (Figure S1A). To further determine the similarities between dSLC5A5 and hSGLT1, we aligned the dSLC5A5 and hSGLT1 amino acid sequences and compared them using UniProt, InterPro, and ExPASy. According to our analyses, the Na^+^-solute symporter family (SSF) signatures are conserved within transmembrane helices 5 and 11 (underlined, Figure S2A) between hSGLT1 and dSLC5A5 and all the Na^+^-binding sites are conserved at positions 77 and 76, 80 and 79, 361 and 389, 364 and 392, and 365 and 393, respectively, between dSLC5A5 and hSGLT1 (red, Figure S2A). Moreover, secondary structure modeling by Protter revealed a similar secondary structure between hSGLT1 and dSLC5A5. There are 14 transmembrane α-helices for hSGLT1 [Figures S2A (yellow highlights) and S2B] and 13 transmembrane α-helices for dSLC5A5 [Figures S2A (yellow highlights) and S2B]. Both the hSGLT1 and dSLC5A5 N-termini face the intestinal lumen while their C-termini face the cellular cytoplasm (Figures S2B and S2B’). Together, these results suggest that dSLC5A5 is the *Drosophila* homolog of SGLT1.

### Whole-body inhibition of dSLC5A5 perturbs systemic glucose homeostasis

We next examined the role of dSLC5A5 in the regulation of systemic glucose homeostasis, a process that is intimately associated with glucose transporter-mediated glucose uptake in the midgut. To determine whether dSLC5A5 loss-of-function affects normal sugar levels in flies, we performed RNA interference (RNAi)-mediated knockdown (KD) of *dSLC5A5* throughout the whole body using the ubiquitous *Actin-Gal4* (*Act-Gal4*), *Daughterless-Gal4* (*Da-Gal4*) or *Ubiquitin-Gal4* (*Ub-Gal4*) drivers and found significantly decreased whole-body glucose level in the *dSLC5A5*-KD flies compared to the RNAi transgene control flies on normal diet (ND) (Figures 1A-1C). The circulating levels of glucose and trehalose were also significantly decreased in the *Act-Gal4*-driven *dSLC5A5*-KD flies relative to the transgene control flies on ND (Figures 1D-1E). Furthermore, food consumption was comparable between the *Act-Gal4*-mediated *dSLC5A5*-KD flies and transgene control flies (Figure 1F), which rules out appetite as a contributing factor in the decreased systemic sugar levels associated with *dSLC5A5* silencing. In contrast, the whole-body KD of *dSLC5A1*, another member of the dSLC5A family, did not significantly affect systemic (Figures S3A-S3C) and circulating (Figures S3D-S3E) sugar levels on ND, pointing to a distinct role of dSLC5A5 in the regulation of normal glucose homeostasis.

**Figure 1.**
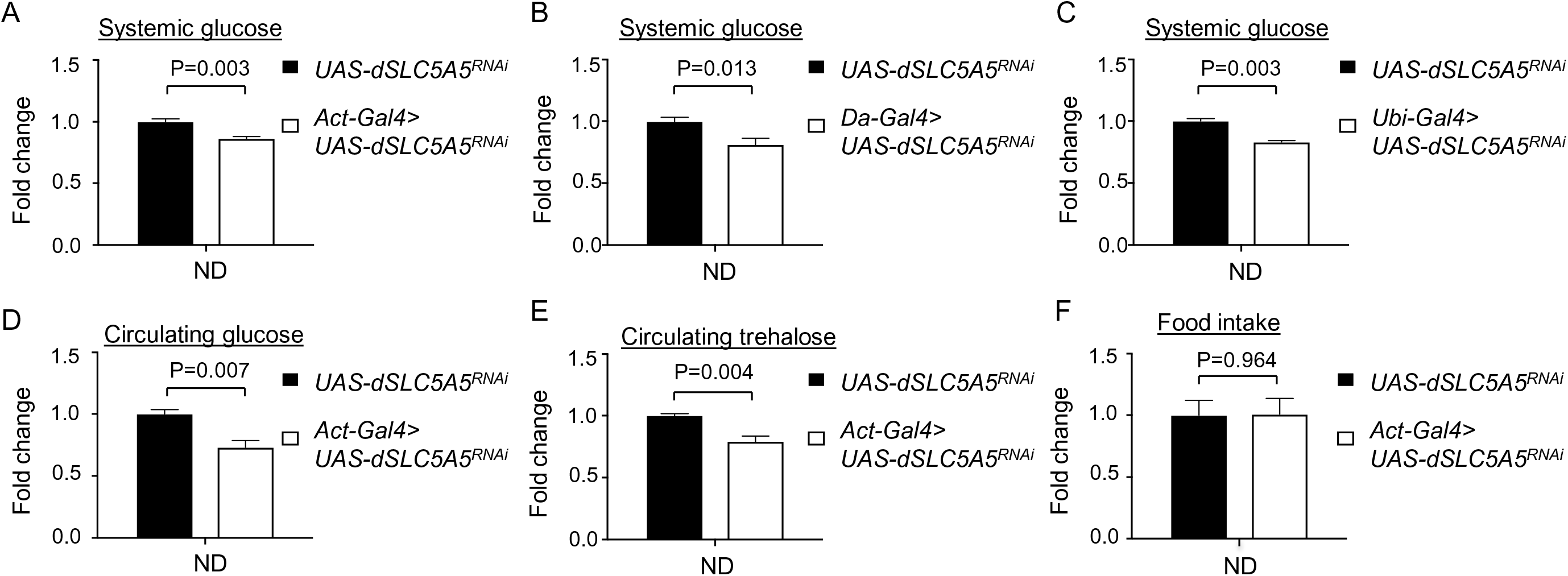
Whole-body knockdown of *dSLC5A5* perturbs systemic glucose metabolism without affecting food intake. (A-C) Systemic glucose levels in 1-week old RNAi transgene control flies (*UAS-dSLC5A5^RNAi^*) or in flies bearing the whole-body KD of *dSLC5A5* using different ubiquitous drivers, *Act-Gal4* (A), *Da-Gal4* (B) or *Ubi-Gal4* (C). Systemic glucose levels (μg/μl) were normalized to whole-body protein (μg/μl). (D, E) Circulating levels of glucose (D) and trehalose (E) in RNAi transgene control flies (*UAS-dSLC5A5^RNAi^*) or in flies with the whole-body KD of *dSLC5A5* mediated by *Act-Gal4* (*Act-Gal4>dSLC5A5^RNAi^*). (F) Food consumption in RNAi transgene control flies (*UAS-dSLC5A5^RNAi^*) or in flies with the whole-body KD of *dSLC5A5* mediated by *Act-Gal4* (*Act-Gal4>dSLC5A5^RNAi^*). Results are expressed as the mean fold change ± SEM from at least five independent experiments. Student’s *t*-test was used to derive *P*-values between the transgene control and KD flies. ND, normal diet.

### dSLC5A5 functions in the midgut to regulate systemic glucose homeostasis

Given that dSLC5A5 is highly expressed in the midgut compared to other tissues (Figure S1A), we asked whether dSLC5A5 acts in the midgut to control systemic glucose homeostasis. To address this, we knocked down *dSLC5A5* specifically in the midgut with two different midgut-specific drivers, *Myo1A-Gal4* and *Caudal-Gal4* (*Cad-Gal4*), each of which resulted in depletion of ∼90% or more of the dSLC5A5 levels in the midgut (Figures S4A-S4B). The midgut-targeted KD of *dSLC5A5* with either driver was associated with significant reduction in systemic glucose levels relative to the RNAi transgene control (Figures 2A-2B). Moreover, circulating glucose (Figure 2C) and trehalose (Figure 2D) levels were similarly decreased to a significant extent in the *Cad-Gal4*-mediated *dSLC5A5*-KD flies compared to the RNAi transgene control flies. In contrast, the KD of *dSLC5A5* specifically in the skeletal muscle did not evoke any obvious changes in systemic glucose level (Figure 2E), circulating glucose (Figure 2F) or circulating trehalose (Figure 2G) levels relative to the RNAi transgene control flies, suggesting that the moderate expression of dSLC5A5 as detected in the skeletal muscles (Figure S1A) does not elicit any obvious effect on normal systemic glucose homeostasis. Furthermore, the targeted KD of *dSLC5A5* in fat body (Figures 2H-2J) or brain (Figures 2K-2M) also did not significantly alter systemic glucose or circulating glucose and trehalose levels compared to the RNAi transgene control flies, which is in agreement with the lack of detectable dSLC5A5 expression in those tissues (Figures S1A). In all, these results reveal an important role of dSLC5A5 in the midgut in maintaining normal systemic glucose homeostasis.

**Figure 2.**
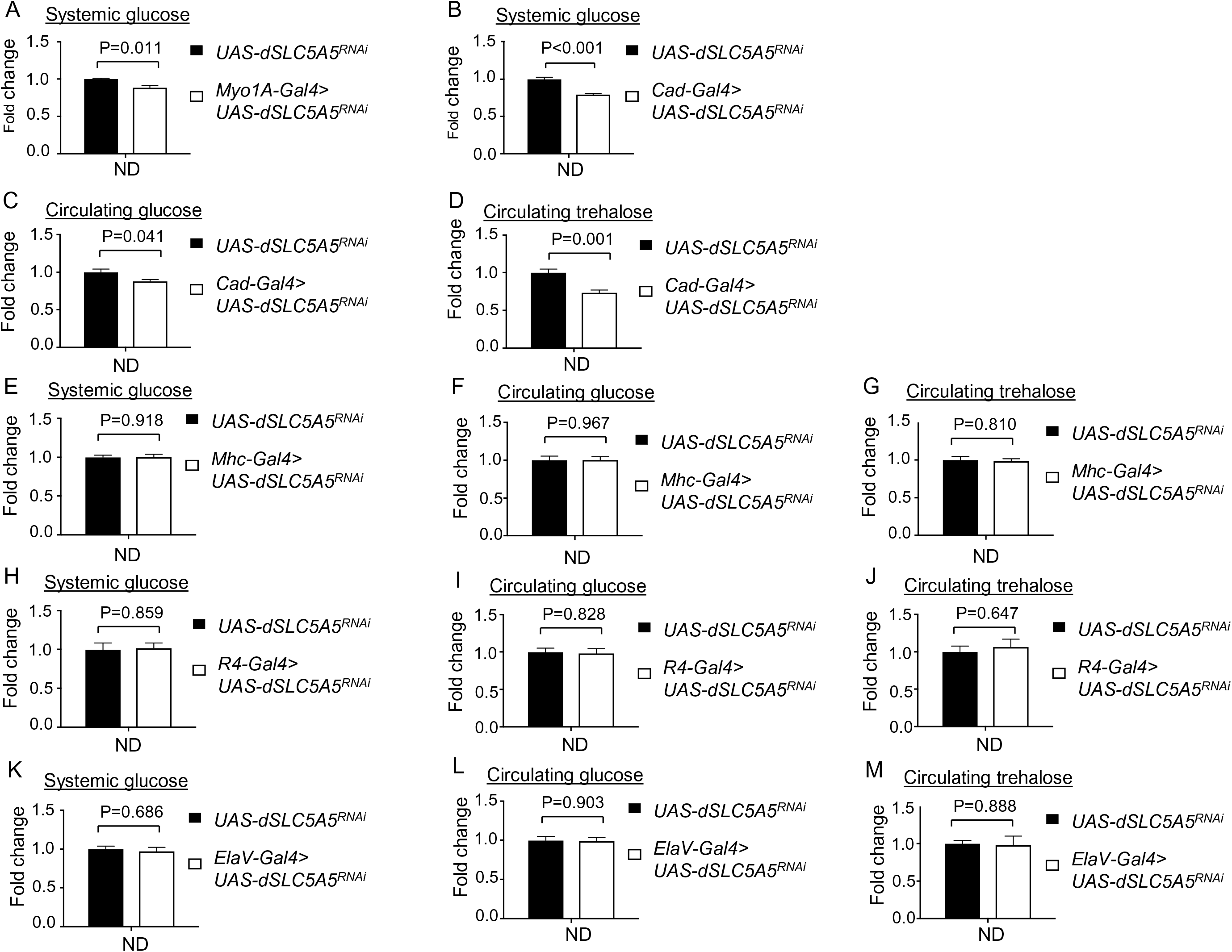
Tissue-specific knockdown of *dSLC5A5* differentially perturbs systemic glucose metabolism. (A-B) Systemic glucose levels in 1-week old RNAi transgene control flies (*UAS-dSLC5A5^RNAi^*) or in midgut-specific *dSLC5A5*-KD flies mediated by *Myo1A-Gal4* (A) or *Cad-Gal4* (B). Systemic glucose levels (μg/μl) were normalized to whole-body protein (μg/μl). (C, D) Circulating glucose levels (C) or circulating trehalose levels (D) in RNAi transgene control flies (*UAS-dSLC5A5^RNAi^*) or in midgut-specific *dSLC5A5*-KD flies mediated by *Cad-Gal4*. (E-G) Systemic glucose levels (E), circulating glucose levels (F), or circulating trehalose levels (G) in 1-week old RNAi transgene control flies (*UAS-dSLC5A5^RNAi^*) or in skeletal muscle-specific *dSLC5A5*-KD flies mediated by *Mhc-Gal4*. (H-J) Systemic glucose levels (H), circulating glucose levels (I), or circulating trehalose levels (J) in1-week old RNAi transgene control flies (*UAS-dSLC5A5^RNAi^*) or in fat body-specific *dSLC5A5*-KD flies mediated by *R4-Gal4*. (L-N) Systemic glucose levels (L), circulating glucose levels (M), or circulating trehalose levels (N) in 1-week old RNAi transgene control flies (*UAS-dSLC5A5^RNAi^*) or in neuronal-specific *dSLC5A5*-KD flies mediated by *ElaV-Gal4*. Systemic glucose levels (μg/μl) were normalized to whole-body protein (μg/μl). Results are expressed as the mean fold change ± SEM from at least five independent experiments. Student’s *t*-test was used to derive *P*-values between the transgene control and KD flies. ND, normal diet.

### Overexpression of dSLC5A5 alters systemic glucose homeostasis

In reciprocal studies, we assessed the gain-of-function of dSLC5A5 on systemic glucose homeostasis. Previous studies have reported fusing various tags including the yellow fluorescent protein to human SGLT1 for the evaluation of SGLT1 abundance in cellular plasma membrane under various conditions (Ghezzi et al., 2017). In the same direction, we also fused a tag (FLAG tag) to the C-terminus of the full-length dSLC5A5 protein for the monitoring of dSLC5A5 abundance and subcellular distribution via FLAG immunostaining. We first crossed the transgenic *UAS-dSLC5A5-FLAG* flies to flies carrying the *Hsp70-Gal4* driver and subjected the progeny flies (*Hsp70-Gal4*>*dSLC5A5-FLAG*) to a transient heat shock, which resulted in an ∼10-fold increase in their whole body *dSLC5A5* mRNA level relative to the *Gal4* driver control flies (Figure 3A). Immunostaining of midguts with an anti-FLAG antibody revealed a stronger FLAG immunosignal in the apical membranes of the enterocytes (marked by the apical membrane marker F-actin) in the *Hsp70-Gal4*>*dSLC5A5-FLAG* flies compared to the *Gal4* driver control flies (Figures 3B-3C’) following heat shock. Quantification of the FLAG immunofluorescence further revealed a near 2-fold increase in dSLC5A5 content in the enterocyte apical membrane in the *Hsp70-Gal4*>*dSLC5A5-FLAG* flies relative to control flies (Figure 3D) following heat shock. Concomitantly, there was a significant elevation in systemic and circulating glucose levels in the *Hsp70-Gal4*>*dSLC5A5-FLAG* flies relative to the *Gal4* driver control flies (Figures 3E-3F) following heat shock. We further targeted the overexpression of dSLC5A5-FLAG in the midgut using *Myo1A-Gal4*, which similarly led to an increased accumulation of dSLC5A5-FLAG in the enterocyte apical membranes (marked by the apical membrane marker F-actin) compared to the transgene control enterocytes (Figure S5A-S5D’). The systemic glucose level in the *Myo1A-Gal4*>*dSLC5A5-FLAG* flies was similarly enhanced compared to the transgene control flies (Figure 3G). These results indicate that dSLC5A5 gain-of-function was associated with glucose metabolic defects that were opposite to those seen with the loss of dSLC5A5 (Figures 1 and 2), which in all corroborate the role of dSLC5A5 as a central regulator of systemic glucose homeostasis.

**Figure 3.**
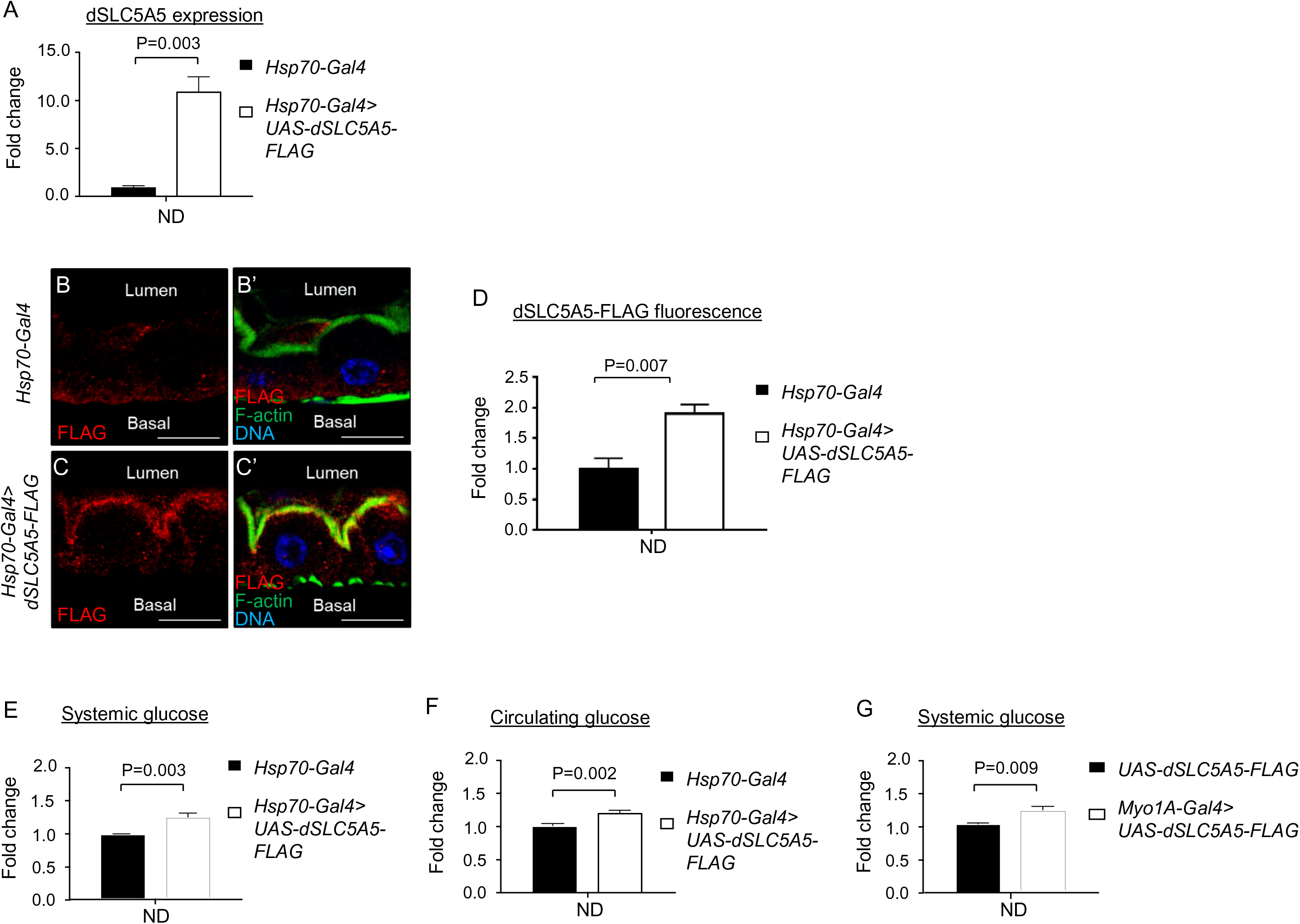
Overexpression of *dSLC5A5* alters systemic glucose metabolism. Quantitative real-time PCR analysis of the whole-fly levels of *dSLC5A5* in *Gal4* control flies (*Hsp70-Gal4*) or in whole-body *dSLC5A5*-overexpressing flies mediated by *Hsp70-Gal4* (*Hsp70-Gal4>dSLC5A5-FLAG*). Flies were subjected to a 1-hour heat shock for induction of dSLC5A5 overexpression. *RPL14* served as an internal control. The data represent the mean fold change ± SEM. Student’s *t*-test was used to derive *P*-values between the control and KD flies. (B-C’) Representative confocal images of the sagittal sections of midguts in *Gal4* control flies (*Hsp70-Gal4*) (B-B’) or in whole-body *dSLC5A5*-overexpressing flies mediated by *Hsp70-Gal4* (C-C’). Flies were subjected to a 1-hour heat shock for induction of dSLC5A5 overexpression followed by midgut isolation and immunostaining for FLAG (red), F-actin (green) and DNA (blue). Scale bars represent 10 μm. (D) Quantification of FLAG immunofluorescence with the mean intensity for the *Hsp70-Gal4>dSLC5A5-FLAG* flies normalized to that of control flies (*Hsp70-Gal4*; set at 1.0). Data represent the mean fold change ± SEM from three independent images. (E-F) Systemic glucose levels in 1-week old transgene control flies *(UAS-dSLC5A5-FLAG)* or in midgut-specific *dSLC5A5*-overexpressing flies mediated by *Myo1A-Gal4 (Myo1A-Gal4>dSLC5A5-FLAG)*. Systemic glucose levels (μg/μl) were normalized to whole-body protein (μg/μl). Results are expressed as the mean fold change ± SEM from at least five independent experiments. Student’s *t*-test was used to derive *P*-values between the transgene control and *Gal4*-mediated RNAi lines. ND, normal diet.

### dSLC5A5 mediates glucose uptake in the midgut

Based on our above observations that an increased accumulation of dSLC5A5 in the apical membranes of the enterocytes (Figures 3B-3C’, S5A-S5D’) was associated with increased systemic and circulating glucose levels (Figures 3E-3G), we posit that dSLC5A5 mediates glucose uptake in the enterocytes. We reasoned that if that is the case, loss of dSLC5A5 will abrogate glucose uptake in the midgut and consequently decreases the levels of glucose in the midgut. Indeed, we detected significantly lowered glucose levels in the *Cad*-and *Myo1A*-mediated *dSLC5A5*-silenced midgut compared to the RNAi transgene control midguts (Figures S6A-S6B, leftmost two bars). We further speculated that in the reciprocal case, gain of dSLC5A5 will augment glucose uptake in the midgut and consequently increase the levels of glucose in the midgut. Indeed, the *Hsp70-Gal4*>*dSLC5A5-FLAG* flies, upon being subjected to a transient heat shock, exhibited significantly heightened glucose content in their midguts compared to midguts in the heat-shocked *Gal4* driver control flies (Figure S6C, leftmost two bars). Together, these results provide evidence supporting a role of dSLC5A5 in regulating glucose absorption in the midgut.

To directly assess glucose uptake function of dSLC5A5, we utilized a non-metabolized fluorescent glucose analog 2-NBDG as an indicator of glucose uptake in the midgut. We first determined whether 2-NBDG uptake by the midgut cells could be inhibited in the presence of glucose. Indeed, in an ex vivo competition assay whereby midguts isolated from starved wild-type *w^1118^* flies were incubated with 300 μM 2-NBDG and increasing concentrations of glucose followed by the analysis of 2-NBDG fluorescence in the enterocytes, we observed a dose-dependent diminution of 2-NBDG fluorescence in the enterocytes upon incubation with increasing amounts of glucose (Figures S7A-S7C). These observations suggest that 2-NBDG and glucose compete for the same glucose transporter to enter the cells, which further indicates that 2-NBDG uptake accurately reflects glucose uptake in the midgut. We then utilized 2-NBDG to evaluate glucose absorption in midguts with genetically altered levels of *dSLC5A5*. In an ex vivo assay, incubating the RNAi transgene control midguts or *dSLC5A5-*silenced midguts with 2-NBDG followed by quantification of 2-NBDG accumulation in the enterocytes revealed a dramatically decreased 2-NBDG fluorescence in the *dSLC5A5*-silenced midguts compared to the RNAi transgene control midguts (Figures 4A-4A’’), with 2-NBDG fluorescence reaching only ∼25% (for the *Cad-Gal4*–mediated *dSLC5A5* silenced midguts) and ∼50% (for the *Myo1A-Gal4*– mediated *dSLC5A5* silenced midguts) of that of the RNAi transgene control midguts (Figure 4B).

**Figure 4.**
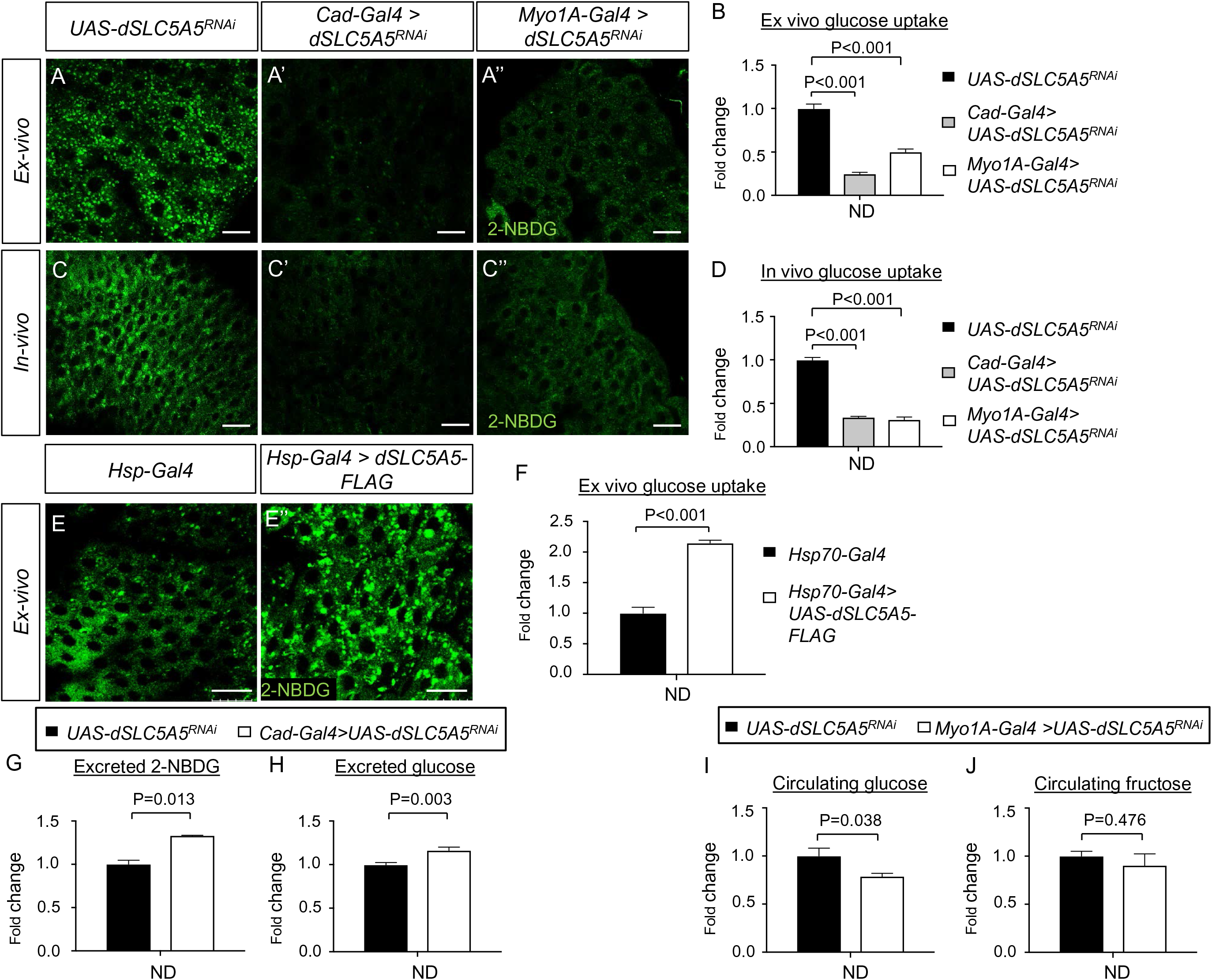
Intestinal-specific inhibition or overexpression of dSLC5A5 disrupts glucose uptake in the enterocytes. (A-C”) Representative confocal images of the intracellular accumulation of 2-NBDG in the *ex-vivo* (A-A”) or *in-vivo* (C-C”) enterocytes of 1-week-old RNAi transgene control flies (*UAS-dSLC5A5^RNAi^*) (A, C) or in midgut-specific *dSLC5A5*-KD flies mediated by *Caudal-Gal4* (A’, C’) or *Myo1A-Gal4* (A”, C”) drivers. Scale bars represent 20 μm. (B, D) Quantification of 2-NBDG fluorescence with the mean intensity for the different genotypes normalized to that of RNAi transgene control flies (*UAS-dSLC5A5^RNAi^*; set at 1.0) in *ex-vivo* enterocyte assay (B) or *in-vivo* enterocyte assay (D). Data represent the mean fold change ± SEM from three independent images. (E-E’) Representative confocal images of the intracellular accumulation of 2-NBDG in the enterocytes of 1-week-old *Gal4* driver control flies (*Hsp70-Gal4*) (E) or in whole-body *dSLC5A5*-overexpressing flies mediated by *Hsp70-Gal4* (E’). Flies were subjected to a 1-hour heat shock for induction of dSLC5A5 overexpression followed by midgut isolation and 2-NBDG incubation *ex-vivo*. Scale bars represent 20 μm. (F) Quantification of 2-NBDG fluorescence with the mean intensity for the *Hsp70-Gal4>dSLC5A5-FLAG* flies normalized to that of control flies (*Hsp70-Gal4*; set at 1.0). Data represent the mean fold change ± SEM from three independent images. (G, H) 2-NBDG levels (G) or glucose levels (H) in the excrement of the RNAi transgene control flies (*UAS-dSLC5A5^RNAi^*) or midgut-specific *dSLC5A5*-KD flies mediated by *Caudal-Gal4* after 2-NBDG or glucose feeding for an hour following overnight starvation. Five 1-week-old female flies per genotype were used for 6-hour excrement collection. (I, J) Circulating glucose levels (I) or circulating fructose levels (J) in RNAi transgene control flies (*UAS-dSLC5A5^RNAi^*) or midgut-specific *dSLC5A5*-KD flies mediated by *Myo1A-Gal4* after glucose or fructose feeding for an hour following overnight starvation. Results are expressed as the mean fold change ± SEM from at least five independent experiments. Student’s *t*-test was used for statistical analysis between the transgene control and *KD* flies. ND, normal diet.

To further corroborate our ex-vivo results, we performed an in vivo assay that entails a short-term (45 minutes) feeding of starved flies with 2-NBDG followed by the rapid isolation of their midguts for visualization and quantification of the 2-NBDG fluorescence intensity. Similar to those observed in the ex vivo assay, 2-NBDG fluorescence was drastically decreased in the *dSLC5A5*-silenced midguts compared to the RNAi transgene control midguts (Figures 4C-4C’’), with 2-NBDG fluorescence reaching only 30-35% (for both the *Cad-Gal4*– and *Myo1A-Gal4*– mediated *dSLC5A5* silenced midguts) of that in the RNAi transgene control midgut (Figure 4D). On the other hand, in the midguts of the *Hsp70-Gal4*–mediated *dSLC5A5-FLAG* overexpressing flies (*Hsp70-Gal4>dSLC5A5-FLAG*) following a transient heat shock, the 2-NBDG fluorescence intensity was increased (up to 2-fold) compared to in the *Gal4* driver control flies (Figures 4E-4E’; 4F). These results indicate that dSLC5A5 positively regulates glucose uptake in the midgut on ND.

Dietary nutrients not absorbed by the enterocytes invariably get expelled from the body into the excreta. We therefore assessed 2-NBDG fluorescence level in the excreta of starved *dSLC5A5-*KD flies and RNAi transgene control flies that were fed transiently with 2-NBDG. A significant increase in 2-NBDG fluorescence level in the excreta of the *Cad-Gal4>dSLC5A5^RNAi^* flies was detected compared to that in control flies (Figure 4G). We then directly quantified the amount of glucose in the excreta and similarly found a significantly higher glucose level in that of the *Cad-Gal4>dSLC5A5^RNAi^* flies relative to that of the RNAi transgene control flies (Figure 4H). In all, these observations indicate an inverse relationship between dSLC5A5-dependent glucose uptake into the midgut enterocytes and glucose expulsion in the excreta, which is consistent with the role of dSLC5A5 acting as a glucose transporter in the midgut.

In the small intestine, SGLT1 is known to transport the monosaccharide glucose across the brush border membrane of enterocytes while fructose, another monosaccharide, is transported via the facilitative diffusion transporter GLUT5 (Koepsell, 2020). To probe whether dSLC5A5 could be involved in fructose transport in the midgut, we fed starved RNAi transgene control flies or flies bearing the midgut-specific KD of *dSLC5A5* (mediated by *MyoA1-Gal4*) with exclusively either glucose or fructose followed by the analysis of circulating glucose or fructose level, respectively. Our results showed that whereas circulating glucose level was decreased significantly in the *MyoA1-Gal4> dSLC5A5^RNAi^* flies relative to RNAi transgene control flies fed with glucose (Figure 4I), the circulating level of fructose remained comparable between both groups of flies fed with fructose (Figure 4J). These results point to a specific role of dSLC5A5 as a glucose transporter in the midgut.

### dSLC5A5 undergoes dynamin-dependent endocytosis in midgut enterocytes

Endocytosis is an important process that controls the amounts of nutrient transporters in the cell surface (Antonescu et al., 2014). One essential protein that is recruited to coated pits for the formation of vesicles during endocytosis is the GTPase dynamin, which was first cloned from the *shibire* temperature-sensitive paralytic *Drosophila* mutant (Grigliatti et al., 1973). Previously, it was reported that overexpression of a dominant-negative form of dynamin in cultured cells did not increase SGLT1 level in the plasma membrane (Ghezzi et al., 2017), suggesting that dynamin does not play a role in the apical endocytosis of SGLT1. However, this finding was obtained from experiments that were performed in a cell line (HEK cells) which is not a physiologically relevant model for human enterocytes and hence might not reflect the endogenous role of dynamin on glucose transporter endocytosis in the intestine. To assess the role of endocytosis in dSLC5A5 abundance and function in the midgut, we abrogated dynamin function by using the *Hsp70-Gal4* to globally knockdown *shibire* and simultaneously overexpress *dSLC5A5-FLAG* (*Hsp70-Gal4>shi^RNAi^; UAS-dSLC5A5-FLAG*). Following a transient heat shock, midguts were isolated and immunostained for FLAG (red; dSLC5A5 amounts) and F-actin (green; enterocyte apical marker). As shown in Figures 5A-5B’, there was a stronger FLAG immunosignal that co-localized with F-actin in the enterocytes of the *Hsp70-Gal4>shi^RNAi^; UAS-dSLC5A5-FLAG* flies compared to the control *Hsp70-Gal4>UAS-dSLC5A5-FLAG* flies. A sagittal (longitudinal) view of the enterocytes similarly revealed an increased abundance of dSLC5A5 in the surface membranes of the enterocytes (marked by F-actin, green) in the *Hsp70-Gal4>shi^RNAi^; UAS-dSLC5A5-FLAG* flies compared to the control *Hsp70-Gal4>UAS-dSLC5A5-FLAG* flies (Figures 5E-5F’). In parallel, there was a reduction in the cytoplasmic level of dSLC5A5 in the enterocytes of the *Hsp70-Gal4>shi^RNAi^; UAS-dSLC5A5-FLAG* flies compared to the control *Hsp70-Gal4>UAS-dSLC5A5-FLAG* flies (Figures 5C-5D’). Quantification of the FLAG immunosignal further indicates a significant 2-fold increase in the dSLC5A5 surface membrane concentration and a significant 0.5-fold reduction in the dSLC5A5 cytoplasmic concentration in the enterocytes of the *Hsp70-Gal4>shi^RNAi^; UAS-dSLC5A5-FLAG* flies compared to the control *Hsp70-Gal4>UAS-dSLC5A5-FLAG* flies (Figures 5G-5G’). These results indicate that dynamin-dependent endocytosis is an important mechanism in regulating the abundance of dSLC5A5 in the apical membrane of the enterocyte.

**Figure 5.**
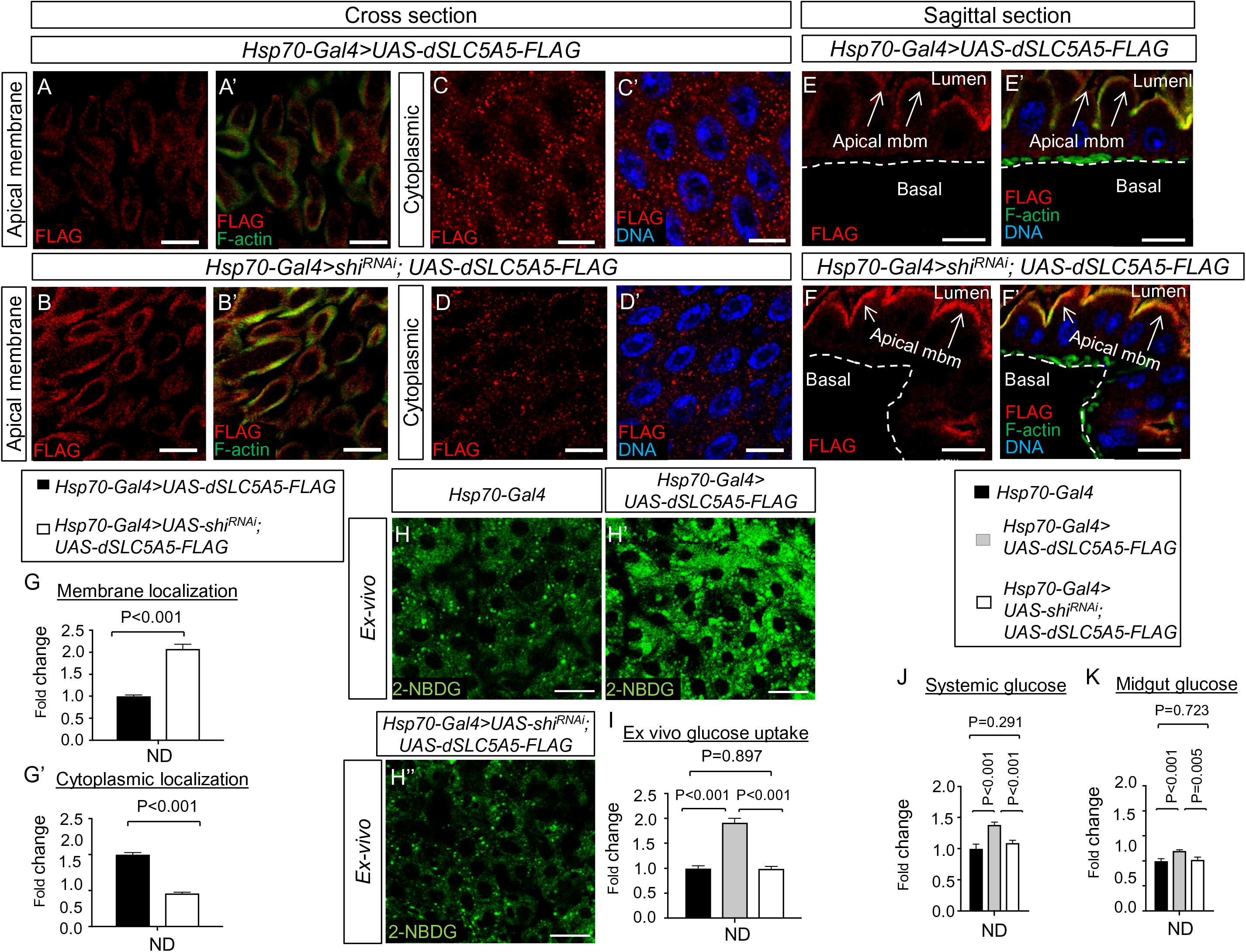
Dynamin-dependent endocytosis regulates dSLC5A5 membrane trafficking and glucose uptake in the enterocytes. (A-F’) Representative confocal images of 1-week-old dSLC5A5-FLAG-overexpressing enterocytes immunostained for FLAG (red), F-actin (green) and DNA (blue) in the absence (A-A’, C-C’, E-E’) or presence (B-B’, D-D’, F-F’) of *shibire*-KD mediated by *Hsp70-Gal4*. Scale bars represent 20 μm. (G-G’) Quantification of the FLAG immunofluorescence in the apical membrane (G) or cytoplasm (G’) of enterocytes with the mean intensity for the *Hsp70-Gal4>shibire^RNAi^*;*dSLC5A5-FLAG* flies normalized to that of control flies (*Hsp70-Gal4*; set at 1.0). Data represent the mean fold change ± SEM from three independent images. (H-H’’) Representative confocal images of the intracellular accumulation of 2-NBDG in the enterocytes of 1-week-old driver control flies (*Hsp70-Gal4*), dSLC5A5-FLAG-overexpressing flies (*Hsp70-Gal4>dSLC5A5-FLAG*), or dSLC5A5-overexpressing and *shibire*-KD flies (*Hsp70-Gal4>shibire^RNAi^*; *dSLC5A5-FLAG*). Flies were subjected to a 1-hour heat shock for induction of dSLC5A5 overexpression followed by midgut isolation and 2-NBDG incubation *ex-vivo*. Scale bars represent 20 μm. (I) Quantification of 2-NBDG fluorescence with the mean intensity for the different genotypes normalized to that of driver control flies (*Hsp70-Gal4*; set at 1.0). Data represent the mean fold change ± SEM from three independent images. (J-K) Systemic glucose levels (J) or midgut glucose levels (K) in 1-week old driver control flies (*Hsp70-Gal4*), dSLC5A5-FLAG-overexpressing flies (*Hsp70-Gal4>dSLC5A5-FLAG*), or dSLC5A5-overexpressing and *shibire*-KD flies (*Hsp70-Gal4>shi^RNAi^*; *dSLC5A5-FLAG*). Systemic glucose levels (μg/μl) were normalized to whole-body protein (μg/μl). Data represent the mean fold change from at least five independent images ± SEM. Student’s *t*-test was used for statistical analysis. ND, normal diet.

To further probe whether dynamin-dependent endocytosis of dSLC5A5 affects dSLC5A5-mediated glucose absorption, we examined the effect of knocking down *shibire* on 2-NBDG uptake in the enterocytes. As expected, there was an increase in 2-NBDG fluorescence intensity (∼1.8-fold) in the midgut of the *Hsp70-Gal4>UAS-dSLC5A5-FLAG* flies relative to the *Hsp70-Gal4* driver control flies following a transient heat shock (Figures 5H-5H’, 5I; see also Figures 4E-4E’). Interestingly, such an increase in 2-NBDG fluorescence intensity in the midgut was suppressed in the *Hsp70-Gal4>shi^RNAi^; UAS-dSLC5A5-FLAG* flies to a level comparable to that of the *Hsp70-Gal4* driver control flies (Figures 5H and 5H’’, 5I). Concomitant with the abrogation of increased glucose uptake into the enterocytes, the heightened systemic (Figure 5J) and intestinal (Figure 5K) glucose levels in the *Hsp70-Gal4>UAS-dSLC5A5-FLAG* flies were normalized in the *Hsp70-Gal4>shi^RNAi^; UAS-dSLC5A5-FLAG* flies on ND. These results indicate that impairment of dynamin-mediated endocytosis suppresses glucose absorption in the enterocytes, despite the entrapping of dSLC5A5 in the enterocyte apical membrane (Figures 5A-5B’; 5E-5F’).

### dSLC5A5 intracellular trafficking is involved in the short-term regulation of dSLC5A5 by glucose

Following endocytosis, SGLT1 can be sorted to the lysosomes for enzymatic proteolysis or recycled back to the plasma membrane in cultured cells (Ghezzi et al., 2017). Furthermore, upon blockade of the lysosomal degradation pathway by proteasome inhibitors (van Kerkhof et al., 2001), SGLT1 content in the plasma membrane of the cultured cells was increased (Ghezzi et al., 2017). To probe whether the same phenomenon also occurs for dSLC5A5 in the enterocytes, we incubated dSLC5A5-FLAG-overexpressing midguts with the proteasome inhibitor MG132 (10 µM) or with PBS for 30 minutes. We observed an enhancement of dSLC5A5-FLAG levels in the apical membrane (marked by F-actin) of the MG132-treated midgut enterocytes relative to PBS-treated midgut enterocytes for both the *Hsp70-Gal4>dSLC5A5-FLAG* (Figures 6A-6B’) and *Cad-Gal4>dSLC5A5-FLAG* (Figures 6F-6G’) flies. Quantification of the apical membrane FLAG immunofluorescence level revealed a nearly 1.5-fold increase in dSLC5A5-FLAG level in the MG132-treated midgut enterocytes than in PBS-treated midgut enterocytes (Figures 6E and 6F). These results support the notion that proteasomal degradation plays a role in the maintenance of dSLC5A5 homeostasis at the apical membrane.

**Figure 6.**
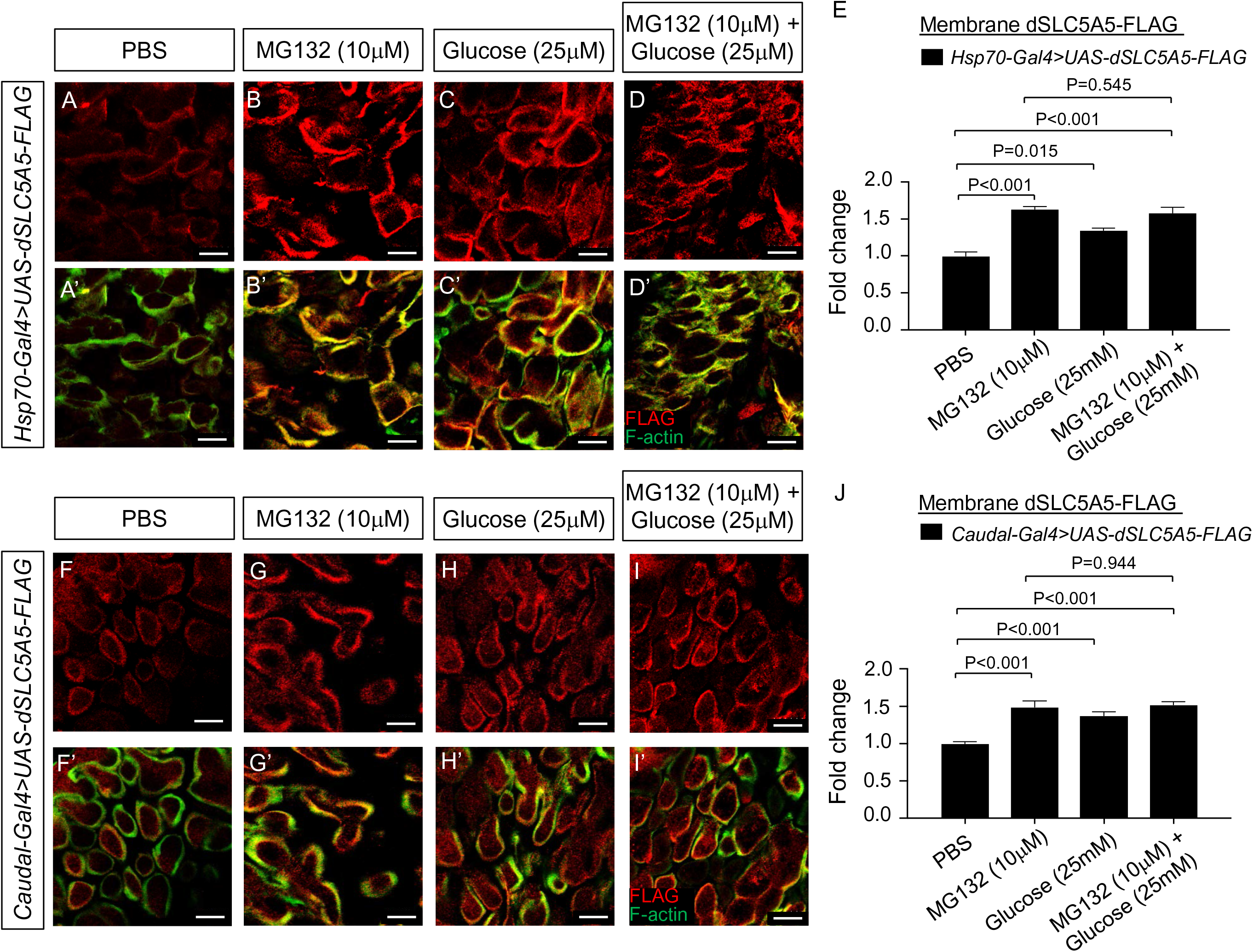
Regulation of dSLC5A5 membrane abundance by proteasome inhibitor, high glucose, or both. (A-D’) Representative confocal images of 1-week-old dSLC5A5-FLAG-overexpressing enterocytes mediated by *Hsp70-Gal4* immunostained for FLAG (red), F-actin (green) and DNA (blue). Flies were subjected to a 1-hour heat shock for induction of dSLC5A5 overexpression followed by midgut isolation and 30 minutes incubation with PBS (A-A’), proteasome inhibitor MG132 (10 µM) (B-B’), high extracellular glucose (25 mM) (C-C’), or in combination (D-D’). Scale bars represent 10 μm. (E) Quantification of the apical membrane FLAG immunofluorescence with the mean intensity for the different genotypes normalized to that of PBS treatment (set at 1.0). Data represent the mean fold change ± SEM from three independent images. (F-I’). Representative confocal images of 1-week-old dSLC5A5-FLAG-overexpressing enterocytes mediated by *Cad-Gal4* immunostained for FLAG (red), F-actin (green) and DNA (blue). Flies were subjected to a 1-hour heat shock for induction of dSLC5A5 overexpression followed by midgut isolation and 30 minutes incubation with PBS (F-F’), proteasome inhibitor MG132 (10 µM) (G-G’), high extracellular glucose (25 mM) (H-H’), or in combination (I-I’). Scale bars represent 10 μm. (J) Quantification of the apical membrane FLAG immunofluorescence with the mean intensity for the different genotypes normalized to that of PBS treatment (set at 1.0). Data represent the mean fold change ± SEM from three independent images.

There has been evidence of a rapid regulation of intestinal SGLT1 by high glucose exposure (Debnam and Sharp, 1993; Debnam et al., 1995; Gorboulev et al., 2012; Sharp et al., 1996), which prompted us to investigate the response of dSLC5A5 in the midgut to acute glucose exposure. When we subjected dSLC5A5-FLAG-overexpressing midguts to high glucose (25 µM) or PBS for 30 minutes, we observed an increased abundance of dSLC5A5-FLAG in the apical membrane (marked by F-actin) of the high glucose-treated midgut enterocytes relative to PBS-treated midgut enterocytes for both the *Hsp70-Gal4>dSLC5A5-FLAG* (Figures 6A-6A’, 6C-6C’) and *Cad-Gal4>dSLC5A5-FLAG* (Figures 6F-6F’, 6H-6H’) flies. Quantification of the apical membrane FLAG immunofluorescence level revealed a significant ∼1.5-fold increase in dSLC5A5-FLAG level in glucose-treated midgut enterocytes than in PBS-treated midgut enterocytes (Figures 6E and 6J). Therefore, like intestinal SGLT1, glucose induces short-term upregulation of dSLC5A5 in the midgut. Interestingly, when we co-incubated dSLC5A5-FLAG-overexpressing midguts with MG132 (10µM) and high glucose (25mM) for 30 minutes, we found no further increase in dSLC5A5-FLAG density in the enterocyte surface membranes compared to either treatment alone, for both the *Hsp70-Gal4>dSLC5A5-FLAG* midguts (Figures 6D-6D’, 6E) and the *Cad-Gal4>dSLC5A5-FLAG* midguts (Figures 6I-6I’, 6J). These observations suggest that dSLC5A5 intracellular trafficking is involved in the short-term regulation of dSLC5A5 by glucose.

### dSLC5A5 is a critical determinant of systemic glucose homeostasis under high sugar diet condition

Our results have revealed an important role of dSLC5A5 in maintaining normal glucose homeostasis (Figures 1-3). We therefore further asked whether dSLC5A5 is also involved in the regulation of glucose homeostasis under high sugar diet (HSD) condition. For that, we subjected newly-eclosed wild-type *w^1118^* flies to a HSD feeding regimen (Figure S8A) that significantly increased their systemic glucose levels and circulating glucose and circulating trehalose levels compared to the ND-fed counterparts (Figures S8B-S8D). The HSD-fed flies also developed insulin resistance, as shown by the reduced phosphorylation of Akt in response to insulin (Figure S8E), as well as obesity, as shown by increased lipid storage in the fat body adipocytes (Figures S8F-S8F’). Together, these observations indicate a dysregulation of glucose metabolism in flies subjected to our HSD feeding regimen. Notably, there was a robust increase in the transcript level of *dSLC5A5* in the midgut (by ∼8-fold, Figure S1B) and in the brain (by ∼3-fold, Figure S1B) compared to the whole body. However, no significant alteration in the transcript level of *dSLC5A5* was detected in the skeletal muscle or fat body relative to whole body (Figure S1B).

To further determine the functional implications of HSD-mediated induction of dSLC5A5, we inhibited whole-body *dSCL5A5* expression under HSD condition using the ubiquitous *Act-*, *Da-*, or *Ub-Gal4* drivers. There was a significant decrease in systemic glucose quantity in the *dSCL5A5*-KD flies compared to the RNAi transgene control flies on HSD (Figures 7A-7C), as well as significantly reduced circulating glucose and circulating trehalose levels in the *Act-Gal4*-mediated dSLC5A5-KD flies compared to RNAi transgene control flies on HSD (Figures 7D-7E). Food intake remained comparable between the KD and control flies on HSD (Figure 7F). In contrast, the whole-body knockdown of *dSLC5A1* did not significantly affect whole-body or circulating sugar content (Figures S3F-S3J) on HSD, which is consistent with its apparent lack of function on systemic glucose homeostasis regulation on ND (Figures S3A-S3E). To further examine if the inhibition of dSLC5A5 specifically in the midgut suppresses HSD-mediated increase in glucose content, we targeted the knockdown of *dSLC5A5* in the midgut with *Myo1A-Gal4* or *Cad-Gal4* and observed a significant reduction in systemic glucose level (Figures 7G-7H), circulating glucose and circulating trehalose levels (Figures 7I-7J), and midgut glucose levels (Figures S6A-S6B, rightmost two bars) relative to the RNAi transgene control flies on HSD. Conversely, overexpression of dSLC5A5 in whole flies (driven by *Hsp70-Gal4* followed by transient heat shock) or specifically in the midgut (using *Myo1A-Gal4*) significantly increased systemic and circulating glucose levels (Figures 7K-7M) and midgut glucose level (Figure S6C, rightmost two bars) relative to the *Hsp70-Gal4* driver control flies on HSD. On the other hand, flies with neuronal-targeted KD of *dSLC5A5* (with *Elav-Gal4*) displayed comparable systemic and circulating glucose and circulating trehalose concentrations with that of the RNAi transgene control flies on HSD (Figures S9A-S9C). Moreover, fat body-specific (with *R4-Gal4*) and muscle-specific (with *Mhc-Gal4*) KD of *dSLC5A5* similarly had no effect on systemic and circulating sugar levels compared to the RNAi transgene control flies on HSD (Figures S9D-S9I). Taken together, these results indicate that dSLC5A5 plays a central role in the midgut in determining HSD-mediated glucose metabolic responses.

**Figure 7.**
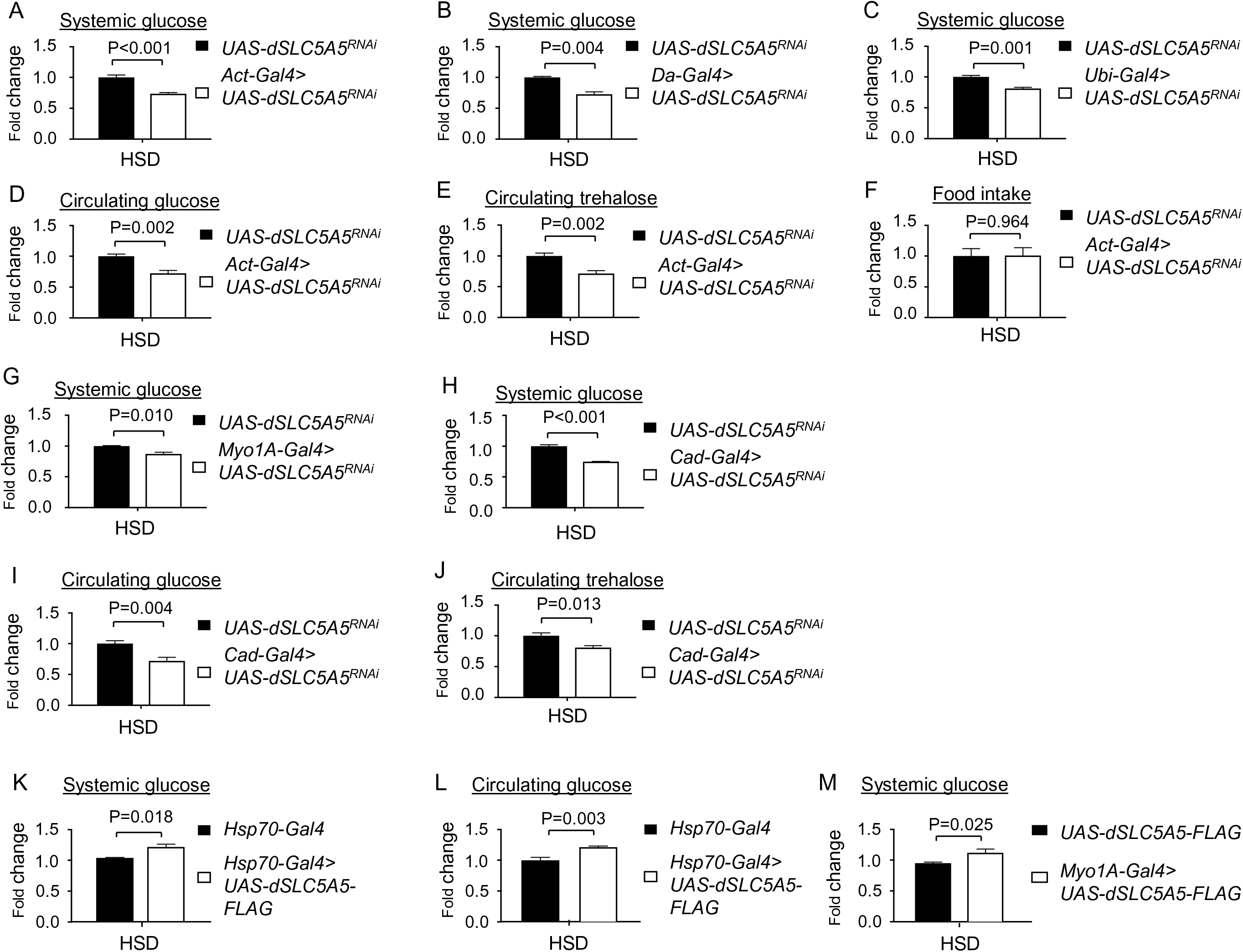
Loss-or gain-of-function of dSLC5A5 alters systemic glucose metabolism on high sugar diet (HSD). (A-C) Systemic glucose levels in 1-week old RNAi transgene control flies (*UAS-dSLC5A5^RNAi^*) or in flies bearing the whole-body KD of *dSLC5A5* using different ubiquitous drivers, *Act-Gal4* (A), *Da-Gal4* (B) or *Ub-Gal4* (C). Systemic glucose levels (μg/μl) were normalized to whole-body protein (μg/μl). (D, E) Circulating levels of glucose (D) and trehalose (E) in RNAi transgene control flies (*UAS-dSLC5A5^RNAi^*) or in flies with the whole-body KD of *dSLC5A5* mediated by *Act-Gal4* (*Act-Gal4>dSLC5A5^RNAi^*). (F) Food consumption in RNAi transgene control flies (*UAS-dSLC5A5^RNAi^*) or in flies with the whole-body KD of *dSLC5A5* mediated by *Act-Gal4* (*Act-Gal4>dSLC5A5^RNAi^*). (G, H) Systemic glucose levels in 1-week old RNAi transgene control flies (*UAS-dSLC5A5^RNAi^*) or in midgut-specific *dSLC5A5*-KD flies mediated by *Myo1A-Gal4* (G) or *Cad-Gal4* (H) on HSD. Systemic glucose levels (μg/μl) were normalized to whole-body protein (μg/μl). (I, J) Circulating glucose levels (I) or circulating trehalose levels (J) in RNAi transgene control flies (*UAS-dSLC5A5^RNAi^*) or in midgut-specific *dSLC5A5*-KD flies mediated by *Cad-Gal4*. (K-M) Systemic glucose levels (K), circulating glucose levels (L), or systemic glucose levels (M) in 1-week old driver control flies (*Hsp70-Gal4*) or dSLC5A5-FLAG-overexpressing flies (*Hsp70-Gal4>dSLC5A5-FLAG*) flies. Systemic glucose levels (μg/μl) were normalized to whole-body protein (μg/μl). Results are expressed as the mean fold change ± SEM from at least five independent experiments. Student’s *t*-test was used to derive *P*-values between the driver control and KD flies. HSD, high sugar diet.

## Discussion

In this study, we report the first identification of a dietary glucose transporter in the *Drosophila* midgut. Our results show that dSLC5A5 is highly expressed in the enterocyte apical membrane and directs glucose uptake which modulates systemic glucose homeostasis. We further show that dSLC5A5 undergoes dynamin-dependent endocytosis in the enterocyte apical membrane. Importantly, we demonstrate that the apical endocytosis of dSLC5A5 contributes to the ability of dSLC5A5 in directing glucose uptake into the enterocytes. Furthermore, our results reveal a short-term upregulation of dSLC5A5 in the midgut by glucose. We provide evidence supporting that glucose upregulates dSLC5A5 abundance in the enterocyte apical membrane via mechanisms that culminate in less dSLC5A5 being directed to the intracellular lysosomal degradation pathway. Finally, we show that dSLC5A5 plays an important role in determining systemic glucose levels on HSD. In all, our studies identify dSLC5A5A5 as the first dietary glucose transporter in the *Drosophila* midgut and uncover new mechanisms that govern the regulation of glucose transporter abundance and activity in the enterocyte apical membranes.

In our work here using the in vivo *Drosophila* midgut system, we demonstrated that that the genetic inhibition of shibire, the *Drosophila* homolog of dynamin, entraps dSLC5A5 in the apical membranes of the enterocytes. On the other hand, a previous study has reported that overexpression of a dominant-negative form of dynamin in cultured cells appeared not to alter the plasma membrane levels of SGLT1 (Ghezza 2017), suggesting that dynamin is not involved in the endocytosis of SGLT1. However, the cultured cell line utilized in thestudy (HEK cells) is not a model for human enterocytes and might not accurately recapitulate the endogenous endocytic regulation of SGLT1. Therefore, the role of dynamin in the endocytosis of glucose transporter in the intestine remains unresolved. While the differing data on the role of dynamin between SGLT1 and dSLC5A5 endocytosis could lie in the different cellular systems utilized and the different approaches used to counter dynamin, our findings provide first evidence supporting an essential role of dynamin on the endocytosis of glucose transporter in a physiological context, which serves as the basis for future studies aimed at elucidating the physiological role of dynamin in the apical endocytosis of SGLT1 in the small intestine.

What is the functional consequence of endocytosis on glucose transporter activity in the enterocytes? Our results showed that inhibition of dSLC5A5 endocytosis was associated with a decrease in glucose uptake into the enterocytes, even though dSLC5A5 abundance in the apical membrane was increased. This interesting finding strongly indicates that endocytosis of dSLC5A5 is required for the ability of dSLC5A5 to transport glucose. It is therefore likely that dSLC5A5 in the apical membrane needs to be refreshed via endocytosis to maintain its glucose uptake ability. The retrieval of the transport-inactive dSLC5A5 from the apical membrane via endocytosis would now allow for newly-synthesized dSLC5A5 to be incorporated into the apical membrane. Alternatively, the endocytosed dSLC5A5 could undergo re-activation in the endosomes before being recycled back to the membrane, in a mechanism that is similar to those involved in the regulation of cell surface receptors such as G protein-coupled receptors (Tan et al., 2004) and cytokine receptors (Stroud et al., 1999). In either case, when endocytosis is abolished, as in our case with the *shibire*-KD enterocytes, the ability of dSLC5A5 to be replenished is correspondingly abrogated, thus culminating in an accumulation of transport-inactive dSLC5A5 in the apical membrane that is paralleled by a decrease in the uptake of glucose in the enterocytes.

Glucose is an important regulator of intestinal glucose transporter expression, as documented by an upregulation in SGLT1 transcription in the jejunum brush border membrane of rats fed on a high carbohydrate diet for several days (Miyamoto et al., 1993), an increase in SGLT1 protein content in the small intestine of mice after glucose gavage (Gorboulev et al., 2012), and an enhancement in glucose uptake in the brush border membrane after rapid exposure of the rat jejunal mucosa to glucose (Debnam and Sharp, 1993; Debnam et al., 1995; Sharp et al., 1996). Our work here further found that dSLC5A5 transcription in the *Drosophila* midgut was also induced in response to a high dietary carbohydrate intake, and that dSLC5A5 protein level in the apical membranes was increased in response to acute high glucose exposure. Together, these data indicate that glucose serves as a critical and conserved regulator of intestinal glucose transporter amounts. The regulation of glucose transporter by glucose appears to critically involve the intracellular trafficking mechanisms of the glucose transporter, including the promotion of the exocytotic pathway of SGLT1 that could contribute to the glucose-induced upregulation of SGLT1 abundance in the small intestine after glucose ingestion (Veyhl-Wichmann et al., 2016). In addition, we observed that in the presence of glucose, the concomitant inhibition of lysosomal degradation by a proteasome inhibitor such as MG132 does not further increase dSLC5A5 level in the enterocyte apical membrane, suggesting that the lysosomal proteolysis of dSLC5A5 minimally contributes to the overall content of membrane dSLC5A5 in the presence of glucose. Potentially, glucose could entrap dSLC5A5 in the apical membrane by eliciting a conformational change in dSLC5A5, or promotes the recycling of dSLC5A5 from the endosomes back to the apical membrane. Either circumstance is likely to lead to decreased amounts of dSLC5A5 being targeted to the lysosomal degradation pathway, thereby lowering the contribution of this intracellular trafficking pathway on the regulation of dSLC5A5 level in the apical membrane. Therefore, abrogation of this pathway in the presence of glucose is expected to minimally impact the level of dSLC5A5 in the enterocyte apical membrane, which was what we observed. The further elucidation of such new mechanisms of intestinal glucose transporter regulation by glucose will contribute towards a greater understanding of SGLT1 regulation in the small intestine.

## Materials and Methods

### Fly Stocks and transgenic flies

All fly stocks were maintained at 25°C on standard medium unless otherwise stated. The following stains were used in this study: *w^1118^* (BDSC BL6326), *Act-Gal4* (BDSC BL3954), *Da-Gal4* (BDSC BL5460), *Caudal* (*Cad*)*-Gal4* (BDSC BL3042), *R4-Gal4* (BDSC BL33832), *ppl-Gal4* (BDSC BL58768), *ElaV-Gal4* (BDSC BL458), *Mhc-Gal4* (BDSC BL38464), *Hsp70-Gal4* (BDSC BL2077), *UAS-dSLC5A5^RNAi^* (BDSC BL63568), *UAS-dSLC5A1^RNAi^* (BDSC BL37273), and *UAS-shibire^RNAi^* (BDSC BL28513). The *Myo1A-Gal4* fly line is a gift from Dr. Bruce Edgar (University of Utah, Salt Lake City). The *dSLC5A5-FLAG* clone (UFO11244) was obtained from the *Drosophila* Genomics Resource Center (DGRC) and contains full-length *dSLC5A5* cDNA with a C-terminus FLAG HA tag in a pUAST vector. *UAS-dSLC5A5-FLAG* transgenic flies were generated by BestGene Inc. (Chino Hills, CA).

### Glucose assay

Whole flies (4 males and 4 females per genotype) or intestines dissected from 12 female flies of each genotype were homogenized in 100 μl PBST (0.05% Triton X-100) with protease inhibitor Cocktail (sigma, P2714). After 10-minute centrifuge (13,000 rps, 4^°^C), 2 μl supernatant was used to detect protein concentration (Bradford assay) and the remaining supernatant was heated for 10 minutes at 70^°^C, followed by 3minutes centrifuged (13,000 rps, 4^°^C). 20 μl from the heated supernatant was diluted in 80μl distilled water (DW) and incubated with 900 μl Glucose Reagent (Thermo Scientific, #TR15421) for 3 minutes at 37^°^C. The absorbance of 340 nm was then measured and the glucose content was calculated with a glucose standard curve. The final glucose concentration (μg/μl) was obtained by normalizing the content to the protein concentration.

### Hemolymph trehalose assay

30-40 one-week old female flies were used for hemolymph collection. Flies punched in the thorax by needles (27G ½”) were transferred to a 0.5 ml microfuge tube that has a small hole at the bottom and filled with a layer of glass wool (Sigma; 20411). The 0.5 ml tube was then placed into a 1.5 ml microfuge tube to be centrifuged at 9,000 g for 5 minutes at 4^°^C. 1 μl hemolymph was carefully removed from the collection tube and diluted in 99 μl trehalase buffer (TB) (5 mM Tris pH 6.6,137 mM NaCl, 2.7 mM KCl) to be immediately heated at 70^°^C for 10 minutes to inactivate the endogenous trehalase. 10 μl out of each heat-treated sample was mixed with 90 μl TB while another 10 μl was added into 90 μl TB with trehalase (TS) [10 μl trehalase (Sigma; T8778-1UN) in 1 ml trehalase buffer] in order to convert trehalose to glucose. After 24-hour incubation at 37^°^C, glucose contents in TB and TS samples were detected with Glucose Reagent (Thermo Scientific, #TR15421). The additional amount of glucose in the TS sample compared to the TB sample was considered to be digested from trehalose.

### Triglyceride assay

Triglyceride assay was performed as previously described (ref). Briefly, adult flies (12 per genotype, 1:1 ratio of males:females) were weighed and homogenized in PBS containing 0.1% Triton X-100 in an amount (μl) that is 8X the total weight of flies (μg) containing protease inhibitor Cocktail (sigma, P2714). The homogenates were centrifuged for 10 min at 13,000 rps (4^°^C) and the supernatants were incubated with Infinity Triglyceride Reagent (Thermo Electron, TR22421) at 37^°^C for 30 minutes. The absorbance of 540 nm was measured and the TG content was calculated from a standard curve constructed with solutions of known TG concentrations (Thermo Electron). The results were normalized to the protein concentration (μg/μl) of each sample (Bradford assay).

### Nile Red staining

Nile Red staining was performed as modified from previously described (ref). Briefly, the fat bodies dissected from adult flies were fixed with 3.7% formaldehyde for 20 minutes at room temperature and then treated with 5 μg/μl of Nile Red solution for 20 minutes. After four times of wash (15 minutes per time) with PBST (0.1% Triton X-100), tissues were mounted in Vectashield media (Vector Laboratories, Inc.) and imaged at 40X magnification using Olympus FV1000 confocal microscope.

### Immunostaining

The midguts were dissected in PBS and fixed with 10% formaldehyde for 10 mins at room temperature. The fixed tissues were then subjected to three rounds of 15 minute-each wash with 0.1% PBST (0.1% Triton X-100 in PBS) and blocked with 5% donkey serum (Jackson ImmunoResearch Laboratories, 107-000-121) for 1 hour at room temperature followed by incubation with primary antibodies in 5% donkey serum at 4°C. The tissues were then washed three times, 10 mins each time in 0.1% PBST, followed by 2 hours incubation with alexa-fluor-conjugated secondary antibodies (Jackson Immunoresearch) in 0.1% PBST at room temperature. After washing as before, tissues were mounted in Vectashield media (Vector Laboratories, Inc.) and viewed under a laser scanning confocal microscope (Olympus FV1000). The following reagents were used for immunostaining: mouse anti-FLAG (1:500, Sigma (F1804)); rat anti-Dilp2 (1:800, a gift from Dr. Pierre Léopold, University of Nice-Sophia Antipolis, France) (Geminard et al., 2009); Alexa Fluor 488 phalloidin (10μM, Invitrogen (A12379)); donkey anti-mouse Cy3 (1:150, Jacson Immunoresearch); donkey anti-rat Cy3 (10μM, Jacson Immunoresearch).

### Heat shock treatment of flies

Flies of the genotype (*Hsp70-Gal4>dSLC5A5-FLAG*) and their respective driver (*Hsp70-Gal4*) and transgene (*dSLC5A5-FLAG*) control flies were incubated at 37^°^C for 40 mins before being subjected to the various assays.

### Short-term ex vivo treatments of midguts

Abdomens of 1-week old female flies were cut opened and each group were incubated with one of the following solutions for 30 mins at room temperature: PBS, PBS with 10uM MG132 (Sigma; M8699), PBS with 25 mM glucose, or PBS with 10 uM MG132 and 25 mM glucose. Intestines were then isolated and fixed in 10% formaldehyde for 10 minutes at room temperature before being incubated with anti-FLAG antibody or Alexa Fluor 488 phalloidin, as described above.

### Ex vivo glucose uptake assay

Following 6-hour starvation with deionized water only, intestines were dissected in ice-cold PBS from 1-week old female flies. A horizontal incision was made to the posterior midgut with the tip of needles (27G ½”) to expose the enterocytes. Samples were incubated with 300 µM 2-NBDG for 45 mins at room temperature. After three brief washes with PBS, tissues were fixed in 3.7% formaldehyde for 15 minutes, followed by three 5-min washes with PBS. Intestines were mounted in Vectashield media.

### In vivo glucose uptake assay

Following 6-hour starvation with deionized water only, 1-week old female flies were fed with 2-NBDG (750 μM in PBS) for 45 minutes at room temperature. Intestines were then dissected in ice-old PBS, fixed in 3.7% formaldehyde for 15 mins, washed in PBS for 3 times (5 minutes per time), and mounted in Vectashield media.

### Excrement glucose assay

Following overnight starvation with only deionized water, 1-week old female flies (N=5 per genotype) were fed with 2-NBDG (250 µM in PBS with blue dye) or ND with 5% blue dye for 1 hour at room temperature. Flies were then transferred into a 1.5 ml Eppendorf tube for excrement collection at room temperature for 6 hours. After discarding the flies, 100 μl of PBS was added into each tube followed by vortexing the tubes to dissolve all the blue-stained excrement deposited in the tube. For 2-NBDG feeding, the absorbance of the blue dye (OD630) and 2-NBDG (488 nm) were measured with the Synergy™ Neo2 Multi-Mode Microplate Reader. For blue ND feeding, 10 µl of the excrement was diluted with PBS (1:20) before blue dye measurement (OD630), and the rest of the excreta (90 μl) was added to 110 µl Glucose Reagent (Thermo Scientific, #TR15421) and incubated for 3 minutes at 37^°^C for glucose level measurement. Final 2-NBDG or glucose content in the excrement was obtained by normalizing the read-out to blue dye amount.

### Glucose feeding assay

One-week old female flies (*n*=30-40 for each genotype) were fed by D-glucose (100 mM in PBS) for 1 hour at room temperature following overnight starvation. Hemolymph (I μl) was collected and diluted in 99 μl PBS and heated at 70°C for 10 minutes. After heating, 20 μl of the diluted mixture was incubated with 980 μl Glucose Reagent (Thermo Scientific, #TR15421) for 3 minutes at 37°C. Readings at 340 nm absorbance were taken and glucose content Cg0 (μg/μl) was calculated using a glucose standard curve. Circulating glucose level equals to 5×Cg0 (μg/μl).

### Fructose feeding assay

One-week old female flies (*n*=30-40 for each genotype) were fed by fructose (20 mg/ml in PBS) for 1 hour at room temperature following overnight starvation. Hemolymph (I μl) was collected and diluted in 99 μl PBS and heated at 70°C for 10 minutes. After heating, 20 μl of the diluted mixture was incubated with 980 μl Glucose Reagent (Thermo Scientific, #TR15421) and 0.2 μl of glucose-6-phosphate isomerase (PGI, Roche, #10127396001), while another 20 μl was incubated with 980 μl Glucose Reagent (Thermo Scientific, #TR15421) only. After 3-minute incubation at 37°C, the glucose level of the mixture with PGI (Cg) or without PGI (Cg0) was calculated based on absorbance of 340 nm using a glucose standard curve. The circulating fructose level (μg/μl) is calculated by ΔCg=Cg-Cg0.

### Real-time quantitative PCR (qPCR)

Total RNA was extracted from tissues in adult flies (n=15 per genotype) using Trizol Reagent (Ambion). cDNA using oligo(dT) were synthesized from total RNA (500ng to 1 μg each) with Superscript III reverse transcriptase (Invitrogen). qPCR amplification reactions were performed in triplicates, by mixing 1 μl of RT product with 10 μl of SYBR qPCR Mastermix (Qiagen) containing the appropriate primers (345 nM of forward primer and 345 nM of reverse primer). Thermal cycling and fluorescence monitoring were performed in a CFX96 (Bio-rad) using the following cycling conditions: 95^°^C for 10 minutes and (95^°^C for 15 seconds, 60^°^C for 1 minute) x 40. Values were normalized with *Rpl14*. Primers used are as follows:

*RpL14* F 5’-TTGCACGTTTCCTCTGTGACG -3’;

R 5’-TCAGAAGGCTCCACCTCCAA -3’;

*dSLC5A5* F: 5’-TCTCTGCCGCTATTGGAGTT -3’

R: 5’-TTCCGTAGGCGTACATTTCC -3’

### Diet feeding regimen

The normal diet (ND) and high sugar diet (HSD) were prepared as described in (Musselman L DMM2011). The feeding regimen using both diets is as follows (see also Figure S7A): Parental flies including w^1118^ flies, RNAi transgene control flies, *Gal4* control flies, or progeny flies from the *Gal4/UAS* crosses were reared on ND until a single uniform of eggs were laid on the food surface (typically in ∼2 days). Parents were then removed and the progenies continued to be reared on the same ND until eclosion. Newly-eclosed male and female adults (equal ratio) were collected and transferred to fresh ND vials or high sugar diet (HSD) vials for rearing for 10 days before being processed for the various analyses.

### Food intake measurement

Food intake for adult flies was measured using the capillary feeder (CAFE) assay as described previously (ref). Briefly, 1-week old adult female flies (N = 10) were weighed and then starved for 16 hours before being transferred to the CAFE chambers comprising an empty food vial with wet cotton at the bottom to maintain humidity. One 100 µL (1 µL/mm) capillary tube with liquid food (400 mM sucrose solution + blue dye) was inserted into each chamber through a cotton plug. Flies were fed at room temperature for 6 hours, after which total liquid food consumption was calculated as the length of colored food at 0 hour minus the length of colored food at 6 hours within the capillary tube (ΔL). A control liquid food chamber without flies was used for the correction of liquid food evaporation (ΔL’). Total food consumption (ΔL-ΔL’) in μl was normalized to weight (g). Food consumption per fly was calculated by further dividing the normalized total food consumption by 10.

### Structural analyses

Sequence alignment and analysis of protein primary structures including prediction of transmembrane helix, putative N-glycosylation sites and Na-binding sites were performed by UniProt (https://www.uniprot.org/), InterPro (https://www.ebi.ac.uk/interpro/) and ExPASy (https://www.expasy.org/) software (Gasteiger et al., 2003; Mitchell et al., 2019; Pundir et al., 2017) with protein sequences in FASTA format extracted from National Center for Biotechnology Information (NCBI). Analyzing tool Protter (http://wlab.ethz.ch/protter/start/) is used to predict secondary structures of proteins, in which only a custom protein sequence in FASTA format or the UniProt protein accession of the protein is needed for the analysis (Omasits et al., 2014).

### Statistical Analysis

All data are presented as the mean ± SEM of the indicated number of replicates. Data were analyzed using the two-tailed Student’s t-test and p<0.05 was considered statistically significant.

## Acknowledgments

We thank our colleagues for thoughtful comments on the manuscript. We are grateful to Dr. Bruce Edgar (University of Utah, Salt Lake City) for sharing the *Myo1A-Gal4* fly line. We thank the Bloomington *Drosophila* stock center and the Vienna *Drosophila* RNAi Center for fly stocks. We thank the Cell Biology Imaging Core Facility at OUHSC for technical assistance on confocal microscopy. This work was supported by grants from the NIH (5R01HL152205-01 to H.-Y.L. and 5R01DK116017-03 to W.W.). H.Y.L. supervised the project, designed most of the experiments, analyzed data, and wrote the paper; Y.L. performed experiments; W.W. designed experiments, analyzed data, and wrote the paper.

## Supplementary Figure Legends

**Figure S1.**
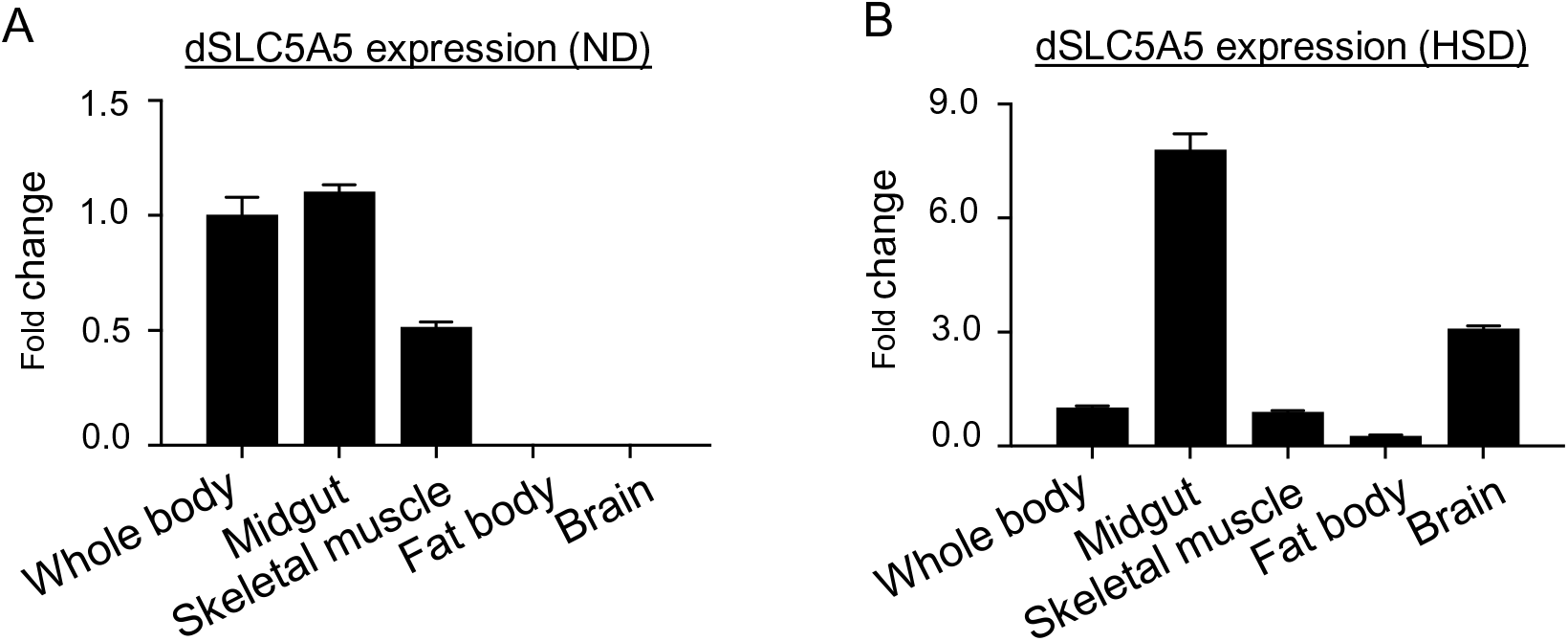
Tissue expression pattern of *dSLC5A5* in *Drosophila* on different diets. (A, B) Quantitative real-time PCR analysis of the transcript levels of *dSLC5A5* normalized to *RPL14* in different fly tissues that were collected from fifteen 1-week-old female *WT* (*w^1118^*) flies reared on normal diet (ND) (A) or high sugar diet (HSD) (B). Whole-body mRNA level of *dSLC5A5* was set as 1. The graphs represent the mean fold change ± SEM.

**Figure S2.**
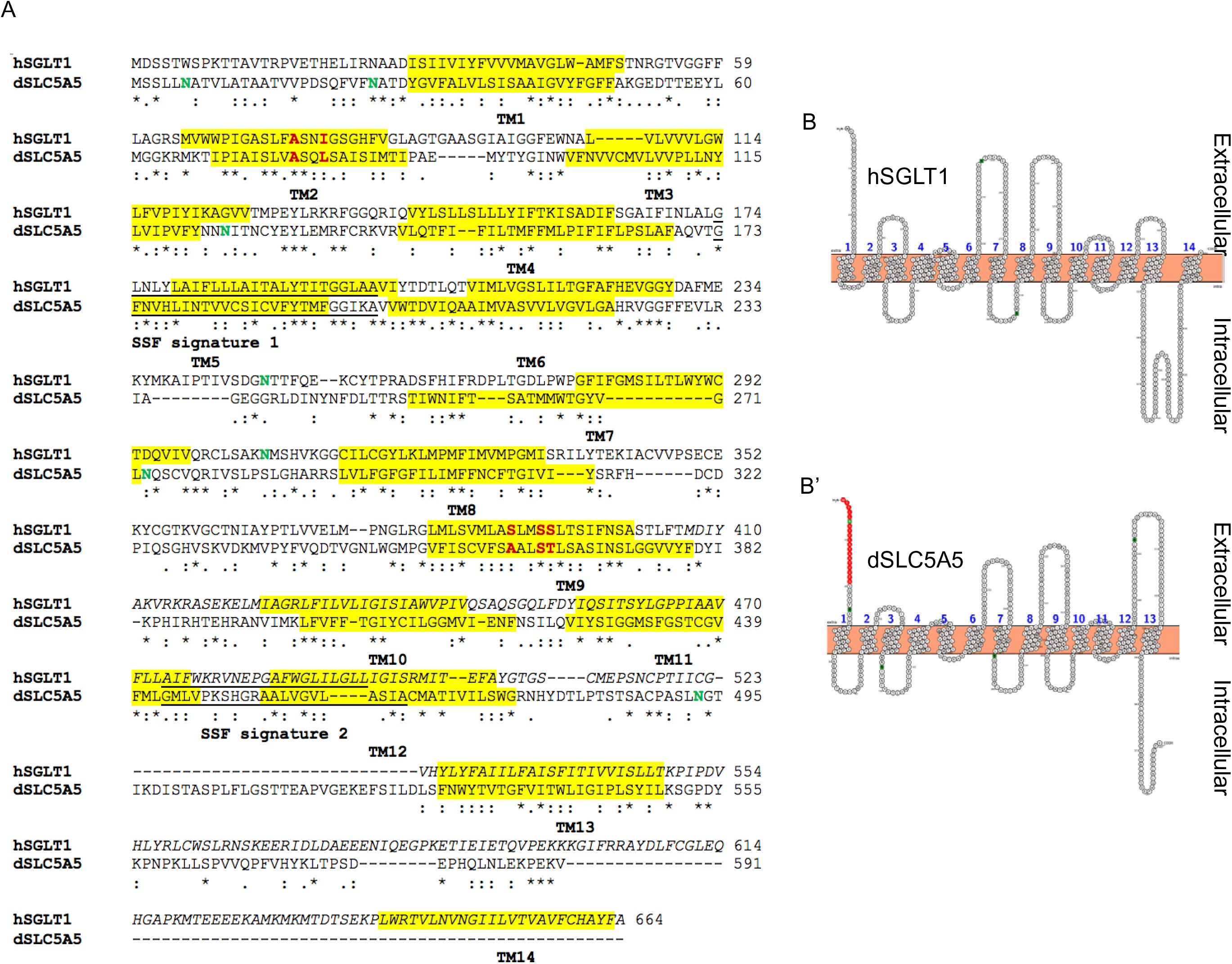
Similarities between hSGLT1 and dSLC5A5. (A) Alignment of amino acid sequences of hSGLT1 and dSLC5A5 produced by UniProt, InterPro and ExPASy. Fully conserved amino acids residues are shown in (*), strongly conserved residues are shown in (:) and weakly conserved residues are shown in (.). Transmembrane regions are shaded in yellow. Sodium: solute symporter family (SSF) signatures are underlined. Putative N-glycosylation sites are in green. Na-binding sites are in red. (B-B’) The secondary structures of hSGLT1 (B) and dSLC5A5 (B’). The signal peptide is highlighted in red. Putative N-glycosylation sites are in green.

**Figure S3.**
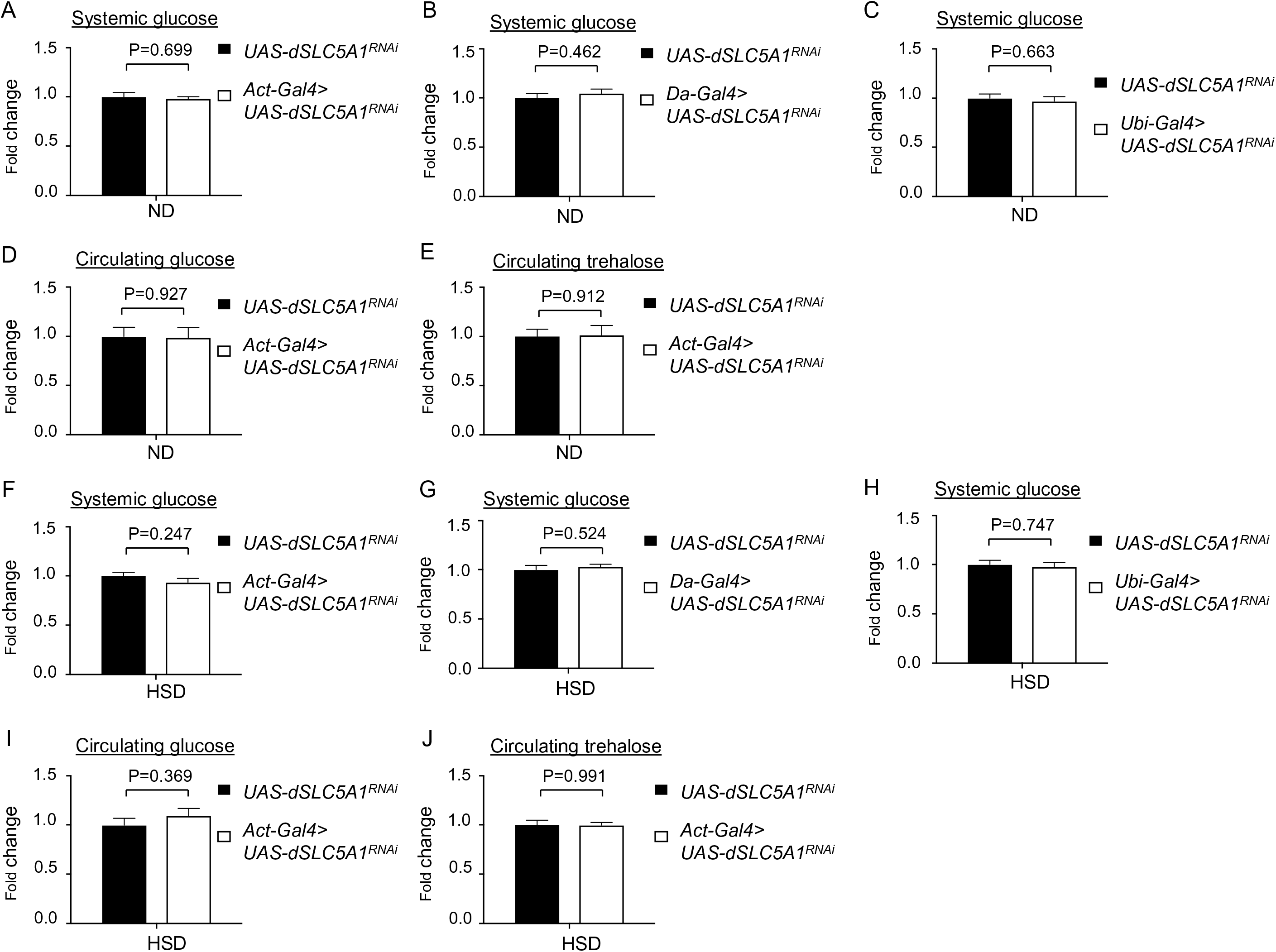
Whole-body down-regulation of *dSLC5A1* does not affect systemic carbohydrate metabolism under different diet conditions. (A-C) Systemic glucose levels in 1-week old RNAi transgene control flies (*UAS-dSLC5A1^RNAi^*) or in whole-body *dSLC5A1*-KD flies mediated by *Act-Gal4* (A), *Da-Gal4* (B) or *Ubi-Gal4* (C) on normal diet (ND). (D, E) Circulating levels of glucose (D) and trehalose (E) in RNAi transgene control flies (*UAS-dSLC5A1^RNAi^*) or in whole-body *dSLC5A1*-KD flies mediated by *Act-Gal4* on ND. (F-H) Systemic glucose levels in 1-week old RNAi transgene control flies (*UAS-dSLC5A1^RNAi^*) or in whole-body *dSLC5A1*-KD flies mediated by *Act-Gal4* (A), *Da-Gal4* (B) or *Ub-Gal4* (C) on high sugar diet (HSD). (I, J) Circulating levels of glucose (D) and trehalose (E) in RNAi transgene control flies (*UAS-dSLC5A1^RNAi^*) or in whole-body *dSLC5A1*-KD flies mediated by *Act-Gal4* on HSD. In A-C and F-H, systemic glucose levels (μg/μl) were normalized to whole-body protein (μg/μl). Results are expressed as the mean fold change ± SEM from at least five independent experiments. Student’s *t*-test was used to derive *P*-values between the transgene control and *Gal4*-mediated RNAi lines on ND or HSD.

**Figure S4.**
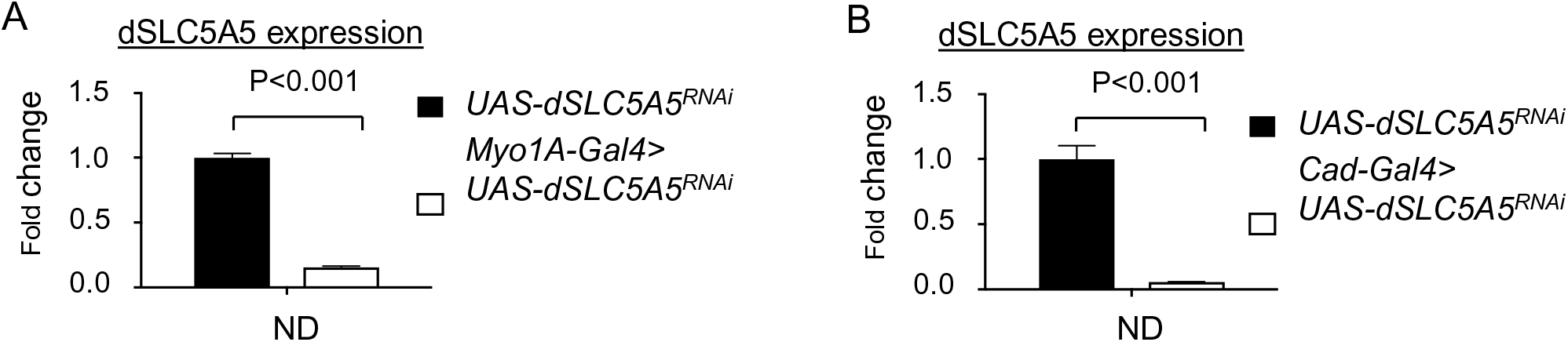
Extent of *dSLC5A5* knockdown mediated by different *Gal4* drivers. (A, B) Quantitative real-time PCR analysis of the midgut transcript levels of *dSLC5A5* in RNAi transgene control flies (*UAS-dSLC5A1^RNAi^*) or in midgut-specific *dSLC5A5*-KD flies induced by *Myo1A-Gal4* (A) or *Caudal-Gal4* (B) on normal diet (ND)*. RPL14* served as an internal control. Midguts were collected from fifteen 1-week-old female flies per genotype for each independent experiment. The data represent the mean fold change ± SEM. Student’s *t*-test was used to derive *P*-values between the control and *Gal4*-mediated KD flies.

**Figure S5.**
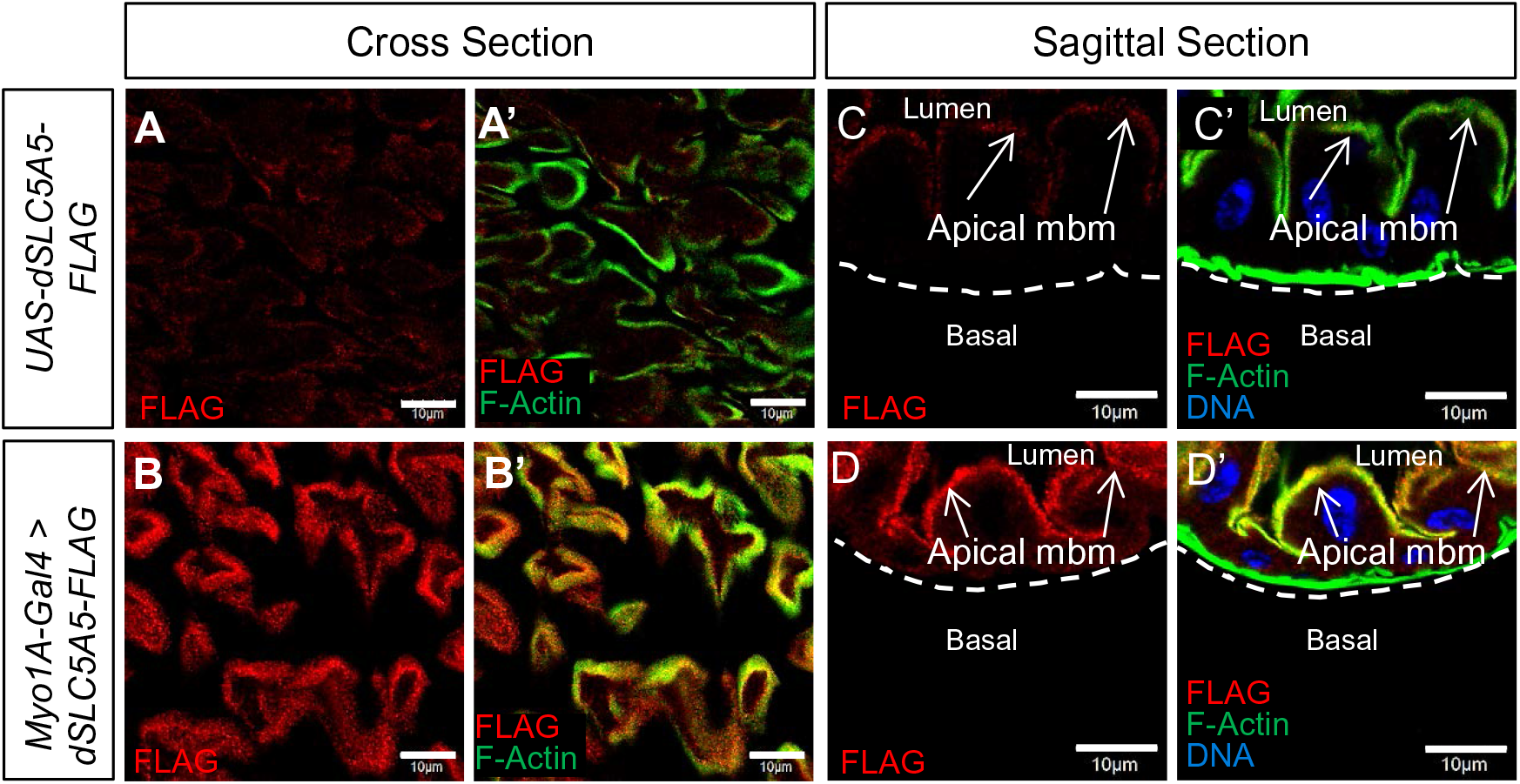
Increased apical membrane abundance of dSLC5A5 in dSLC5A5-overexpressing midgut enterocytes. (A-B’) Representative confocal images of the cross sections of midguts in the transgene control flies (*UAS-dSLC5A5-FLAG*) (A-A’) or in midgut-specific *dSLC5A5*-overexpressing flies mediated by *Myo1A-Gal4* (B-B’). (C-D’) Representative confocal images of the sagittal sections of midguts in the transgene control flies (*UAS-dSLC5A5-FLAG*) (C-C’) or in midgut-specific *dSLC5A5*-overexpressing flies mediated by *Myo1A-Gal4* (D-D’). In all cases, midguts were dissected in 1-week old flies and immunostained for FLAG (red), F-actin (green) and DNA (blue). Scale bars represent 10 μm.

**Figure S6.**
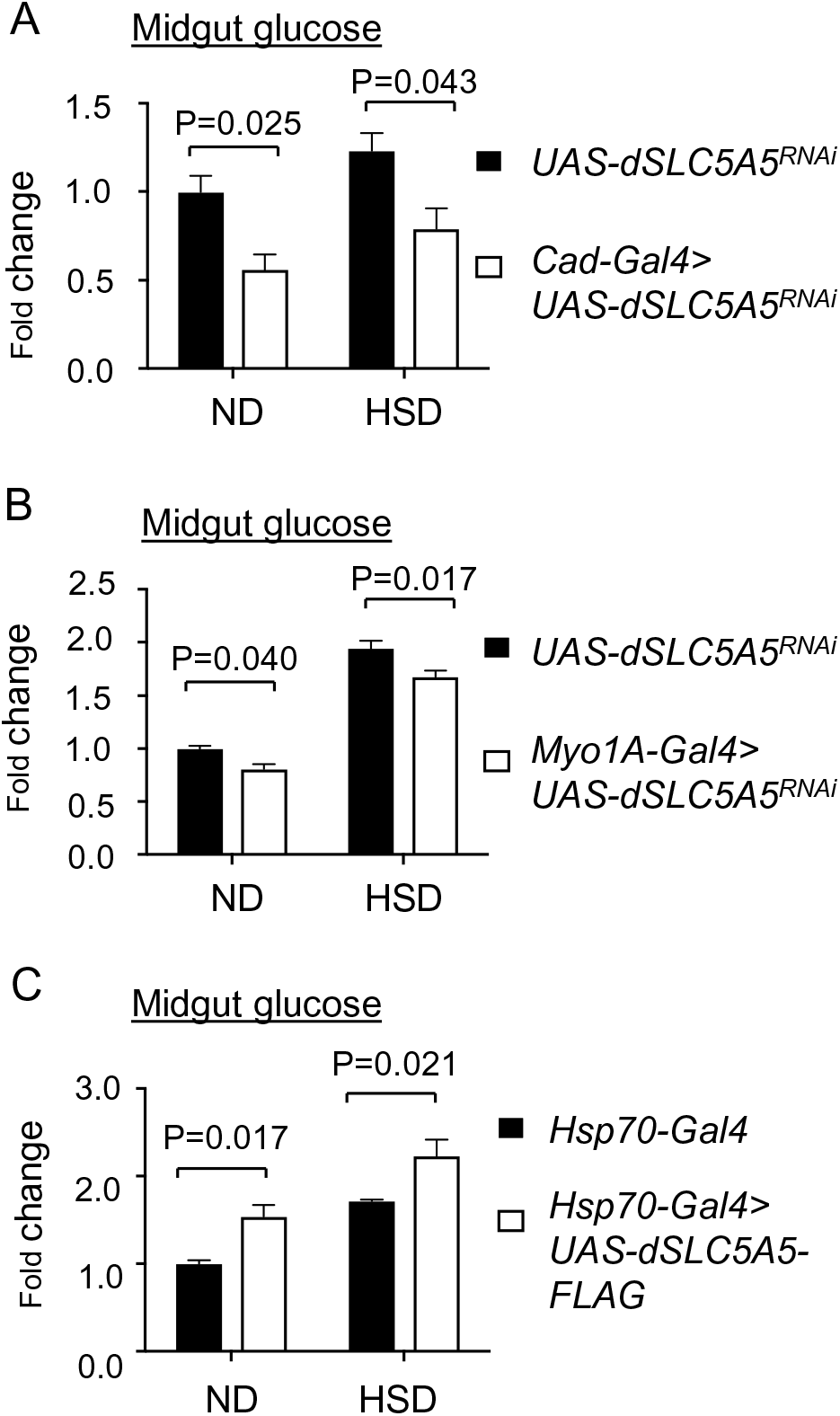
Midgut-specific knockdown or overexpression of dSLC5A1 decreases or increases, respectively, midgut glucose levels. (A-B) Midgut glucose levels in 1-week old RNAi transgene control flies (*UAS-dSLC5A1^RNAi^*) or in midgut-specific *dSLC5A1*-KD flies induced by *Caudal-Gal4* (*Cad-Gal4>dSLC5A1^RNAi^*) (A) or *Myo1A-Gal4* (*Myo1A-Gal4>dSLC5A1^RNAi^*) (B) on normal diet (ND) or high sugar diet (HSD). (C) Midgut glucose levels in 1-week old *Gal4* driver control flies (*Hsp70-Gal4*) or in whole body-*dSLC5A1-FLAG* overexpressing flies mediated by *Hsp70-Gal4* (*Hsp70-Gal4>dSLC5A1-FLAG*) on ND or HSD. Midgut glucose levels (μg/μl) were normalized to midgut protein levels (μg/μl). Results are expressed as the mean ± SEM from at least five biological replicates. Student’s *t*-test was used for statistical analysis between the each control and KD-or overexpressing flies under ND or HSD condition.

**Figure S7.**
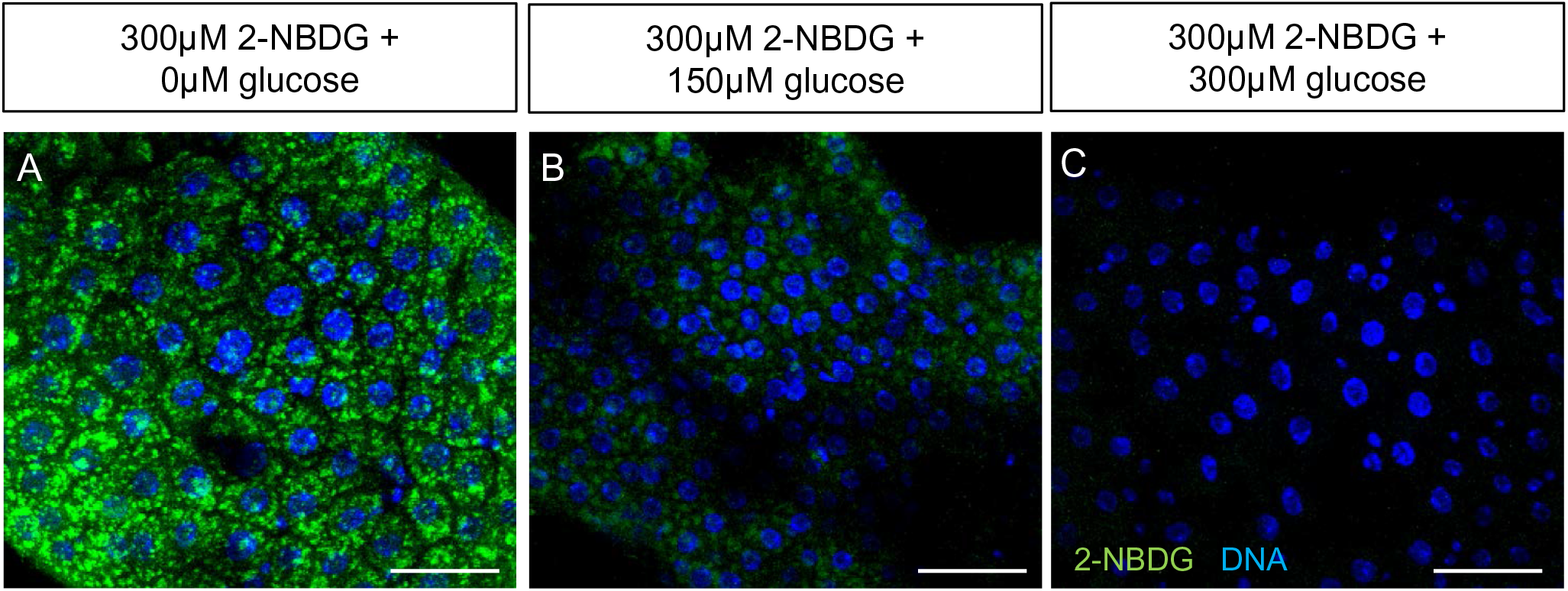
2-NBDG and glucose are absorbed through the glucose transporter in midgut enterocytes. (A-C) Representative confocal images of midguts dissected from 4 day-old female *w^1118^* flies and incubated with 300 μM 2-NBDG together with 0 μM (A), 150 μM (B), and 300 μM (C) of glucose for 30 minutes at room temperature following a 30-minute starvation in PBS. Scale bars represent 30 μm.

**Figure S8.**
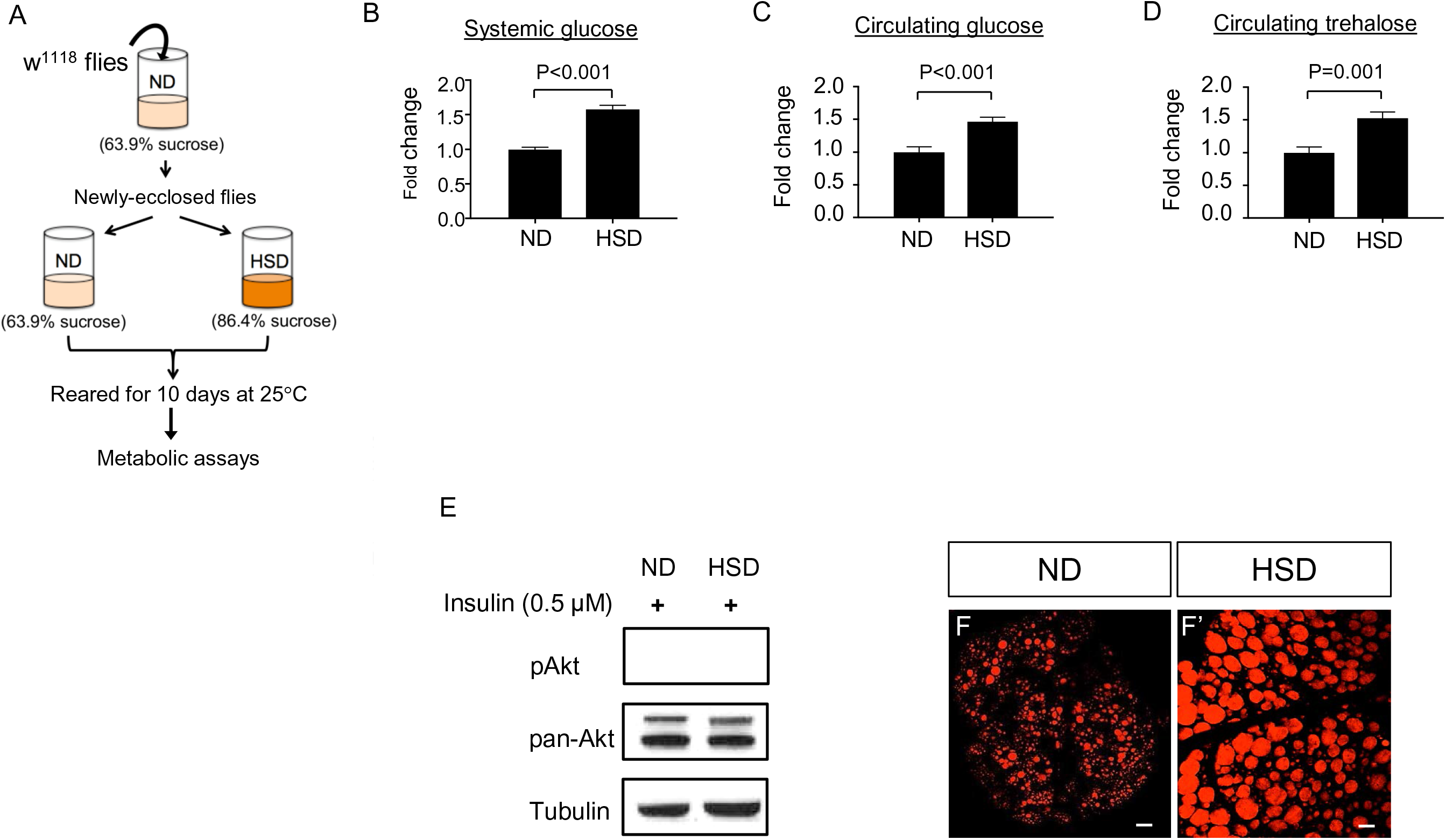
High sugar diet (HSD) feeding of *Drosophila* generates metabolic disorders. (A) Schematic workflow of the high sugar diet (HSD) feeding regimen of wild-type *w^1118^* flies. (B-D) Levels of whole-body glucose (B), circulating glucose (C), and circulating trehalose (D) in *w^1118^* flies after 10-day feeding on normal diet (ND) or HSD post-eclosion. Systemic glucose levels (μg/μl) were normalized to total protein of whole flies (μg/μl). Results are expressed as the mean ± SEM from at least five biological replicates. Student’s *t*-test was used for *P*-value analysis. (E) Western blot analysis of phosphorylated Akt (p-Akt) level in 1-week old *w^1118^* flies on ND or HSD after 15-minute incubation with 0.5 μM human insulin. (F-F’) Representative confocal images of lipid droplets stained by Nile Red in the fat body of *w^1118^* flies under ND (F) or HSD condition (F’). Scale bars represent 20 μm.

**Figure S9.**
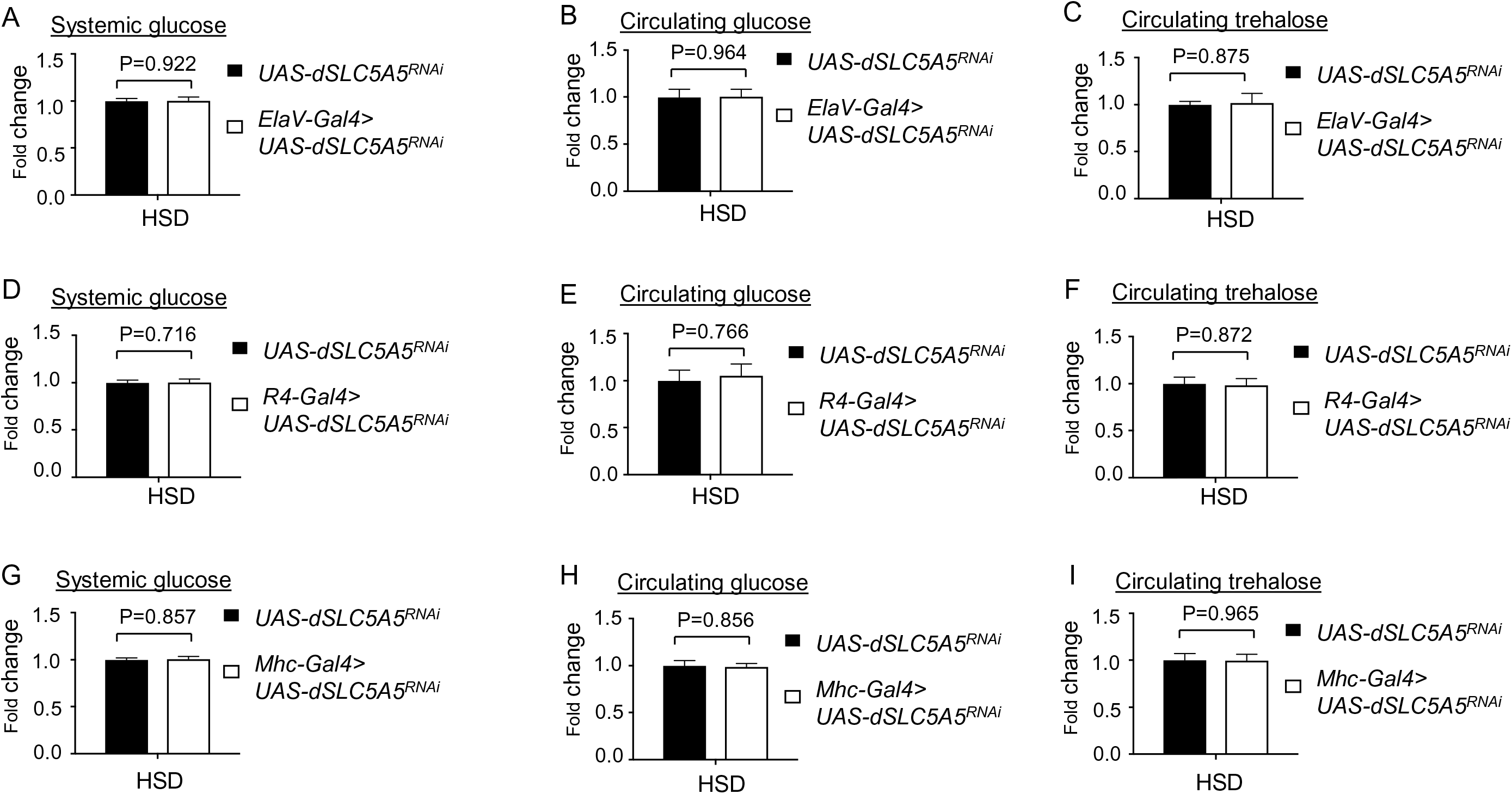
dSLC5A5 inhibition in the brain, fat body, or skeletal muscle does not affect systemic and circulating glucose levels under high sugar diet (HSD) condition. (A, D, G) Systemic glucose levels in 1-week old RNAi transgene control flies (*UAS-dSLC5A5^RNAi^*) or flies bearing the neuronal KD of *dSLC5A5* using *ElaV-Gal4* (A), the fat body KD of *dSLC5A5* using *R4-Gal4* (D), or the skeletal muscle KD of *dSLC5A5* using *Mhc-Gal4* (G) on HSD. (B, E, H) Circulating glucose levels in 1-week old RNAi transgene control flies (*UAS-dSLC5A5^RNAi^*), flies bearing the neuronal KD of *dSLC5A5* using *ElaV-Gal4* (B), flies bearing the fat body KD of *dSLC5A5* using *R4-Gal4* (E), or flies bearing the skeletal muscle KD of *dSLC5A5* using *Mhc-Gal4* (H), on HSD. (C, F, I) Circulating trehalose levels in 1-week old RNAi transgene control flies (*UAS-dSLC5A5^RNAi^*) or flies bearing the neuronal KD of *dSLC5A5* using *ElaV-Gal4* (C), flies bearing the fat body KD of *dSLC5A5* using *R4-Gal4* (F), or flies bearing the skeletal muscle KD of *dSLC5A5* using *Mhc-Gal4* (I), on HSD. Systemic glucose levels (μg/μl) were normalized to whole-body protein (μg/μl) in A, D, and G. Results are expressed as the mean fold change ± SEM from at least five independent experiments. Student’s *t*-test was used to derive *P*-values between the transgene control and *Gal4*-mediated RNAi lines on HSD.

## Notes

### Competing Interest Statement

The authors have declared no competing interest.

